# Human CYP1A2 metabolism of organophosphorus pesticides and nerve agent surrogates

**DOI:** 10.64898/2026.10.20.760143

**Authors:** Pratik Shriwas, Matthew Politeski, Brian Scarpitti, Andre Revnew, Thomas R. Lane, Sean Ekins, Christopher M. Hadad, Craig A. McElroy

## Abstract

CYP1A2 contributes 13–15% of hepatic CYP450 content and plays a critical role in the metabolism of approximately 10% of pharmaceutical drugs in the human body. CYP1A2 has been previously reported to bioactivate and detoxify both the thion (P=S) and oxon (P=O) forms of organophosphorus (OP) compounds. In this study, we determined the capacity of CYP1A2 to degrade OP compounds using LC-MS/MS, as well as the inhibition of its activity by OP compounds using the previously developed pFluor50 assay. CYP1A2 was selective in metabolizing thions over oxons, and several OP compounds inhibited its activity in a time- and dose-dependent manner. To further investigate the mechanism of CYP1A2–OP interactions, molecular docking studies were performed using a crystal structure of CYP1A2 (PDB ID: 2HI4). Linear correlation analysis between CYP1A2 inhibition at 10 µM with 30 minute preincubation and the % OP compound remaining at 2 hours (R = -0.71, P<0.001) showed a moderate but significant correlation. Based on these findings, CYP1A2 was identified as a potential bioscavenger for OP compound metabolism, supporting the future integration of experimental and *in silico* data for developing artificial intelligence (AI) / machine learning (ML) models for prediction of CYP1A2-mediated metabolism of existing and emerging OP compounds, including those of toxicological and chemical warfare relevance.

## Introduction

Organophosphorus (OP) pesticides act as covalent inhibitors of human acetylcholinesterase (AChE) in erythrocytes and the neurosynaptic cleft, thereby causing accumulation of the neurotransmitter acetylcholine and disrupting cholinergic transmission (Aroniadou-Anderjaska *et al*., 2023; Chen *et al*., 2023). This activity makes OP compounds dangerous to human health, leading to acute as well as long-term neurological disorders (Rendic and Guengerich, 2021). OP poisoning remains a major global health concern, with ∼10–20% fatality rates per incident (Eddleston *et al*., 2008), and an estimated 258,000–371,000 deaths worldwide each year (Gunnell *et al*., 2007). In agricultural settings, pesticide exposure remains widespread, with studies reporting that up to 44% of farmers experience symptoms consistent with pesticide poisoning (Damalas and Koutroubas, 2016). Likewise, quantifiable pesticide residues were found in approximately 41% of the 110,829 food samples analyzed in the European Union in 2022 (European Food Safety Authority (EFSA) *et al*., 2024), which can ultimately contribute to neurotoxicity and cancer (Alavanja, Hoppin and Kamel, 2004).

Recently, several high-profile individual and mass-casualty incidents have involved the use of another highly toxic class of OP compounds, chemical warfare nerve agents (CWNAs). These include the 1995 sarin attack in the Tokyo subway (Okumura *et al*., 1996), the sarin attack on Khan Shaykhun in Idlib Province, Syria, in April 2017, in which approximately 90 civilians were killed and several hundred more were injured (OPCW, 2017), and the poisoning of Alexei Navalny in Russia in 2020 with a highly potent Novichok agent (Haslam *et al*., 2022). Overall, the acute and chronic toxicity induced by OP pesticides and the fatalities caused by CWNAs indicate an urgent need to counter OP-mediated toxicity by investigating its mechanisms in depth.

OP compounds can be grouped into two chemically distinct subclasses, thions (P=S) and oxons (P=O), depending on which element is double-bonded to the phosphorus atom (Bharate *et al*., 2010). This distinction is critical with respect to CYP450-mediated metabolism, as thions are often desulfurized to the active oxon form by various CYP450 enzymes in the human body, whereas CWNAs, being oxons, cause a rapid onset of AChE inhibition and subsequent fatality (Thompson, Prins and George, 2010). Currently, clinical treatment of OP poisoning relies on anticholinergic agents such as atropine and on oxime-based AChE reactivators (Eddleston *et al*., 2008; Kharel *et al*., 2020). However, the broader use of these treatments is limited by pharmacokinetic constraints in the central nervous system. Quinone methide precursor compounds are an emerging class of AChE reactivators that overcome these limitations, but they have not yet entered clinical trials and still require further testing in *in vivo* models (Yoder *et al*., 2017; Zhuang *et al*., 2018; Clay *et al*., 2024; Lovins *et al*., 2024, 2026). At present, there is no medication that acts as a complete and effective medical countermeasure to OP poisoning. One complementary strategy is the use of bioscavengers, enzymes that bind or degrade OP compounds before they reach their targets. Butyrylcholinesterase, which binds OP compounds stoichiometrically, and paraoxonase 1, which hydrolyzes only a limited range of OP compounds, have been investigated for this purpose, so enzymes that can degrade structurally diverse OP compounds catalytically remain of interest.

Recently, our group and others identified OP pesticides and CWNA surrogates that were degraded by human liver microsomes (HLMs) using NMR and targeted LC-MS/MS metabolomics (Agarwal *et al*., 2023). However, because HLMs contain many enzymes, such as CYP450 isoforms, amidases, and esterases, among others, the results were not conclusive with respect to identifying the individual proteins that metabolized these toxic compounds.

Cytochrome P450 enzymes provide an essential defensive system against xenobiotics by carrying out Phase I oxidative metabolism of drugs, exogenous chemicals, and endogenous substrates (Elfaki *et al*., 2018; Zuo *et al*., 2022). Among these enzymes, CYP1A2 contributes to ∼10% of clinical drug metabolism while constituting 13–15% of hepatic CYP450 content (Klein *et al*., 2010; Zanger and Schwab, 2013). CYP1A2 has also previously been found to metabolize various pesticides such as azinphos-methyl, chlorpyrifos, diazinon, parathion, methyl parathion and malathion at concentrations representative of human exposure (Tang *et al*., 2001; Buratti *et al*., 2003, 2005; Ellison *et al*., 2012; Rendic and Guengerich, 2015). Critically, CYP1A2 had two opposing mechanisms in metabolism, desulfurization of thions such as chlorpyrifos to the AChE- inhibiting bioactive oxon form and the other one involving methyl parathion in dearylation leading to detoxification (Tang *et al*., 2001; Ellison *et al*., 2012). Because these studies did not investigate parent compound metabolism, it becomes important to test this using LC-MS/MS. Because CYP1A2 has a narrower spectrum of drug targets and exhibits greater selectivity than previously tested CYP450s such as CYP2C9 (Shriwas, Noonchester, Revnew, *et al*., 2026), CYP2C19 (Vignaux *et al*., 2023) and CYP3A4 (Shriwas, Noonchester, Scarpitti, *et al*., 2026), we tested its ability to metabolize OP compounds and CWNA surrogates in this study. In our earlier work, many of the OP compounds tested were not metabolized by CYP2C9, CYP2C19, or CYP3A4, which prompted us to examine an isoform with a different substrate profile.

This study investigates the capacity of recombinant CYP1A2 to metabolize structurally diverse OP pesticides and CWNA surrogates using targeted LC-MS/MS analysis. We hypothesized that CYP1A2 would degrade a subset of OP compounds, that time-dependent inhibition measured with the high-throughput pFluor50 assay would reflect this turnover, and that docking parameters would help explain both. CWNA surrogates, which mimic the structures of CWNAs but are less hazardous to handle, were included for experimental work, whereas the authentic CWNAs were examined only by docking. Thirteen of 22 OP pesticides were metabolized by more than 50% after 2 hours, as was the CWNA cyclosarin surrogate CMP after 1 hour, which was unexpected given the selectivity of CYP1A2 for its substrates. Interestingly, oxons and thions had different inhibitory activities, with more thions showing significant time-dependent inhibition at 10 µM and more oxons doing so at 1 µM, further indicating the selectivity of CYP1A2 between these two classes. Overall, we found that the experimental results from inhibition at 10 µM OP compound with a 30-minute preincubation and LC-MS/MS-detected degradation were correlated across all tested OP pesticides at 2 hours (R = -0.71, P<0.001) and 1 hour (R = -0.60, P<0.01).

Further, the *in silico* molecular docking parameter binding efficiency at the shortest reactive distance (BEeff (SD)) correlated with the time-dependent inhibition at 10 µM (R = 0.36, P<0.05). 3D docking studies revealed that cyclosarin and its surrogate CMP adopt a similar docking pose. These experimental findings, along with *in silico* data, can be used to develop predictive AI/ML models of CYP1A2-mediated metabolism of OP pesticides as well as of the more toxic CWNA surrogates and the CWNAs themselves.

## Materials and Methods Chemicals

All organophosphorus (OP) pesticides used in this study were obtained as neat compounds as described previously (Shriwas et al., 2026) and were utilized without additional purification. CWNA surrogates were synthesized in the Department of Chemistry and Biochemistry at The Ohio State University. The chemical structures of all OP compounds examined in this work, presented in either oxon or thion form, are shown in Figures S1 and S2 (Supplementary Figures). Compounds containing geometric (E/Z) or stereochemical (R/S) isomers were evaluated as mixed isomeric preparations. Key physicochemical and toxicological properties of the OP compounds—including hydrogen bond donor and acceptor counts, molecular weight, log P, polar surface area, number of rotatable bonds, and reported LD₅₀ values—are summarized in Supplementary Table S1.

## pFluor50 CYP1A2 Inhibition Assay

CypExpress™ 1A2 (Sigma, MTOXCE1A2), containing full-length human recombinant CYP1A2, recombinant human P450 NADPH oxidoreductase, and a proprietary mix of components; magnesium chloride (MgCl2; M9272) and reduced nicotinamide adenine dinucleotide phosphate (NADPH tetrasodium salt; 481973-Millipore) were purchased from Sigma Aldrich (St. Louis, MO). The fluorogenic substrate 7-ethoxy-3H-phenoxazin-3-one or resorufin ethoxy ether (Eres) for CYP1A2 (# 16122) was purchased from Cayman Chemical (Ann Arbor, MI), while α-naphthoflavone (#N5757) was obtained from Sigma Aldrich (St. Louis, MO). All 384-well Opti-Plates, 0.1 M potassium phosphate buffer, and OP stock solutions were prepared as described previously (Shriwas, Noonchester, Revnew, *et al*., 2026). A 28.8 µL aliquot of enzyme-containing master mix comprising 2 mg/mL CYP1A2, 0.52 mM NADPH, and 2.64 mM MgCl2 was mixed with 1.2 µL of OP stock (0.5 or 0.05 mM), and the mixture was preincubated for either 10 or 30 minutes. After the incubation period was complete, 30 µL of 4 µM Eres was added to each well. Resorufin formation was then measured by fluorescence (excitation 560 nm, emission 590 nm) for 10 minutes using a BioTek Synergy H1 plate reader with parameters as previously described (Shriwas *et al*., 2025). α-Naphthoflavone was used as a positive control for inhibition of CYP1A2. Slope normalization was performed as previously described (Shriwas, Noonchester, Revnew, *et al*., 2026) using Equation 1 below.

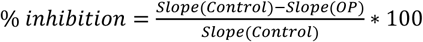

Each experiment was performed using at least three technical replicates, and data analysis was performed using Prism 10.4.2 software (GraphPad Software Inc.). Pairwise comparisons using the two-tailed unpaired t-test (with the Holm-Šídák correction for multiple comparisons) and a significance threshold of 0.05 were used for statistical analysis.

## LC-MS/MS Metabolism Assays

*CYP1A2 Clearance:* BioDuro (Shanghai, China) performed the CYP1A2 LC-MS/MS metabolism study. The entire protocol was followed as previously described (Shriwas, Noonchester, Revnew, *et al*., 2026). Briefly, OP compounds (1 µM final reaction concentration) were pre-incubated with recombinant 15 pmol CYP1A2 at 37 °C for 5 minutes before addition of 1 mM NADPH. 10% of the reaction was separated and subsequently quenched with a mixture of 5 ng/ml terfenadine and 10 ng/ml tolbutamide solution in acetonitrile at 0, 5, 30, 60 and 120 minutes. Surrogate measurements were taken at 0, 5, 15, 30, and 60 minutes. Phenacetin was used as a positive control CYP1A2 substrate. Parameters were calculated as follows:

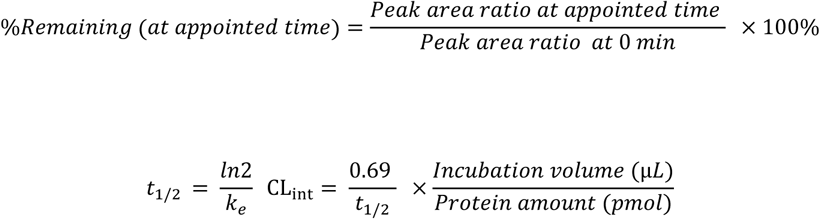

where ke is the elimination/degradation rate constant, and CLint is the intrinsic clearance rate.

## Molecular Docking Analysis

The crystal structure of human cytochrome P450 CYP1A2 with bound alpha-naphthoflavone (PDB 2HI4) was used for subsequent molecular docking simulations (Chemical Computing Group (CCG) | accessed, 04-09-2026). The structure was imported into the Molecular Operating Environment (MOE) software (v2024.06), where chain A was isolated, and α-naphthoflavone was removed (Sansen *et al*., 2007). Protein preparation was performed using MOE’s QuickPrep function, which applies homology modeling to complete missing residues, adds hydrogens, determines protonation states, and refines the structure using the Amber14:EHT force field (Labute, 2009; Maier *et al*., 2015). Several residues at the termini of chain A were missing.

Therefore, they were capped by acylating the N-terminus and amidating the C-terminus using MOE’s protein builder. The receptor was then converted to pdbqt format using the utility tool prepare_receptor in the ADFR suite (Ravindranath *et al*., 2015). For the ligands, the OP pesticides’ SMILES strings were imported into MOE and prepared using the Wash method, where hydrogens were added, and the 3D coordinates were reconstructed. An energy minimization was performed with the Amber14:EHT force field (Maier *et al*., 2015). The prepared compound structures were then converted to pdbqt format using the Python program Meeko (v0.5.0), which supports atom typing and enhanced sampling of macrocycles during docking (Forli and Botta, 2007).

## Docking Method

Molecular docking calculations were conducted using AutoDock Vina version 1.1.5 with the grid space centered at 4.95, 24.78, and 24.36 Å and expanded by 18, 20, and 20 points in the x-, y-, and z-directions, respectively, to encompass the heme iron and nearby residues. The protein was kept rigid during the docking calculations. Before docking the OP pesticides, alpha- naphthoflavone was re-docked onto its crystallographic binding site to validate the protocol.

Subsequently, each OP pesticide was docked into the receptor at an exhaustiveness level of 200, generating up to 20 unique poses.

## Correlation and Regression Analyses

Docking parameters such as shortest distance and binding energy were correlated with the experimental data from the LC-MS/MS metabolism and pFluor50 inhibition assays as described previously (Shriwas, Noonchester, Revnew, *et al*., 2026). Correlation analysis was also performed between the molecular parameters to investigate relationships among them. Multiple linear regression analysis overfitted the CYP1A2 data, so individual linear regression models were used instead. The correlation analyses used the inhibition values at 1 and 10 µM after 10 and 30 minutes, the time-dependent inhibition (the difference in percent inhibition between the 10- and 30-minute preincubations at each concentration), and the percentage of OP remaining by LC-MS/MS at 1 or 2 hours; these were correlated with the molecular parameters, docking-derived binding energies, and reactive distances. Results of the correlation analyses are presented as the Pearson or Spearman correlation coefficient (R) and significance value (P), along with the equation of the regression line.

## Results

### CYP1A2 is inhibited by oxons and thions in a time-dependent manner

CYP1A2 activity was quantified using the previously developed pFluor50 fluorogenic method with Eres as a substrate (Shriwas *et al*., 2025). In this study, the same protocol was used to investigate the activation or inhibition of 47 OP compounds on CYP1A2 activity. α- Naphthoflavone was used as a positive control for CYP1A2 inhibition (Figure 1 A-B). We observed striking differences in time-dependent inhibition by thions and oxons at 1 and 10 µM. Fourteen of 21 thions and 15 of 26 oxons showed time-dependent variation in inhibition at 10 µM. However, 24 of 26 oxons but only 14 of 21 thions showed time-dependent variation in inhibition at 1 µM (Figure 1A-B). The overall summary of the variations at 1 and 10 µM between 10 and 30 minutes is presented in a heatmap in Figure 1C. Many OP pesticides such as CFVP, EPP, FMP, IBF, MCP and NEDPA were inhibitors at 1 µM after 10 minutes but became activators after 30 minutes. However, apart from MPF and NEDPA at 10 µM after 10 minutes, no OP compounds acted as activators at either the 10- or 30-minute time points at this concentration. Many OP compounds at 10 µM, such as CPO, CFA, and CTP, among others, showed increased inhibition at 30 minutes, whereas their inhibition at 1 µM decreased between 10 and 30 minutes. These time-dependent variations in CYP1A2 activity suggest mechanism-based inhibition or increasingly tight binding and highlight how concentration can alter the way OP compounds interact with CYP1A2.

**Figure 1.**
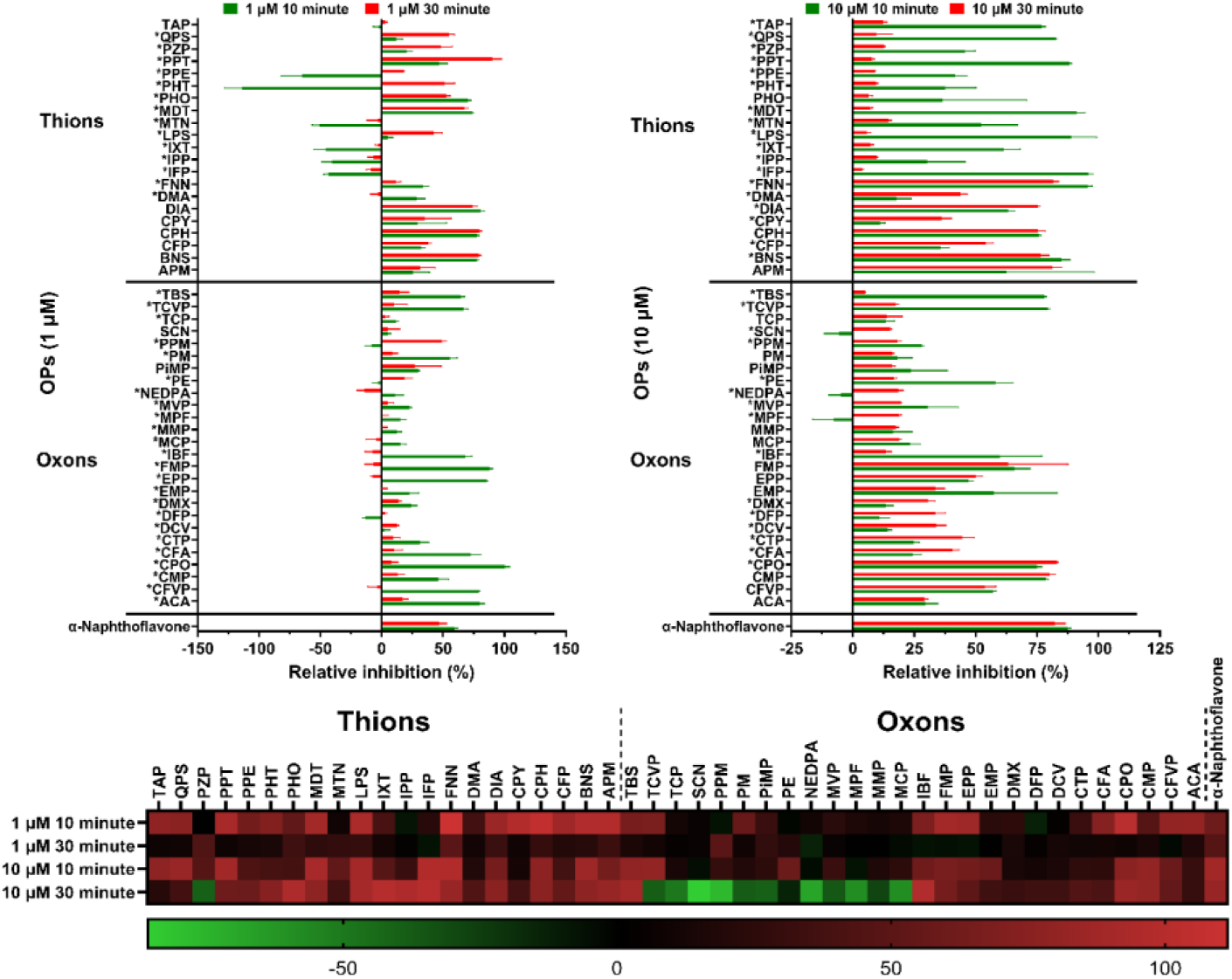
Relative inhibition of CYP1A2 by OP compounds as measured using the pFluor50 fluorogenic assay. **A**. Relative inhibition of CYP1A2 activity by thions and oxons at 1 µM concentration with 10 minute (green bars) or 30 minute (red bars) preincubation. B. Relative inhibition of CYP1A2 activity by thions and oxons at 10 µM concentration with 10 (green bars) or 30 (red bars) minute preincubation. C. Heatmap showing variations in CYP1A2 activity due to OP inhibition at 1 and 10 µM after 10 or 30 minute preincubation. * Represents OP compounds wherein the inhibition of CYP1A2 changes significantly between 10 and 30 minute preincubation. α-Naphthoflavone was used as a positive control for inhibition of CYP1A2.

### Thions are more susceptible to CYP1A2-mediated metabolism than oxons

LC-MS/MS experiments were performed with CYP1A2 to directly determine the degradation of OP compounds (Table 1). OP quantification was performed at various time points up to 1 hour (CWNA surrogates) or 2 hours (all other compounds tested). Phenacetin was used as a positive control. We found that all thions but one were degraded by more than 50%, with MTN metabolized by only ∼30% after 2 hours. QPS and FMN were metabolized almost completely within 120 minutes of incubation. IPP, MDT, IXT, TAP, PHO, PHT, PPT, and TEM were degraded by 50–80% within 2 hours (Figure 2B). Among the oxons, only FMP was degraded by more than 80% and EPP by ∼65% after 2 hours, with TCVP, CFVP, and TCP metabolized by around 25–40% (Figure 2A). MVP, PPM, DCV, ACA and PFN were not degraded after 2 hours (Figure 2A). Among the surrogates, only CMP (a cyclosarin surrogate) was degraded, by ∼60%, whereas PiMP (a soman surrogate), EMP (a VX surrogate), and NEDPA (a tabun surrogate) were not metabolized at all during the 1-hour exposure, suggesting that CYP1A2 was selective towards specific surrogates (Figure 2C). Overall, thions had a lower mean percentage of parent compound remaining (43.62%) than oxons (78.85%) after 1 hour of CYP1A2 incubation, with a significant mean difference of −35.24 ± 9.89 (mean ± SE; P = 0.0016; 95% CI, −55.65 to −14.83) (Figure 2D).

**Figure 2.**
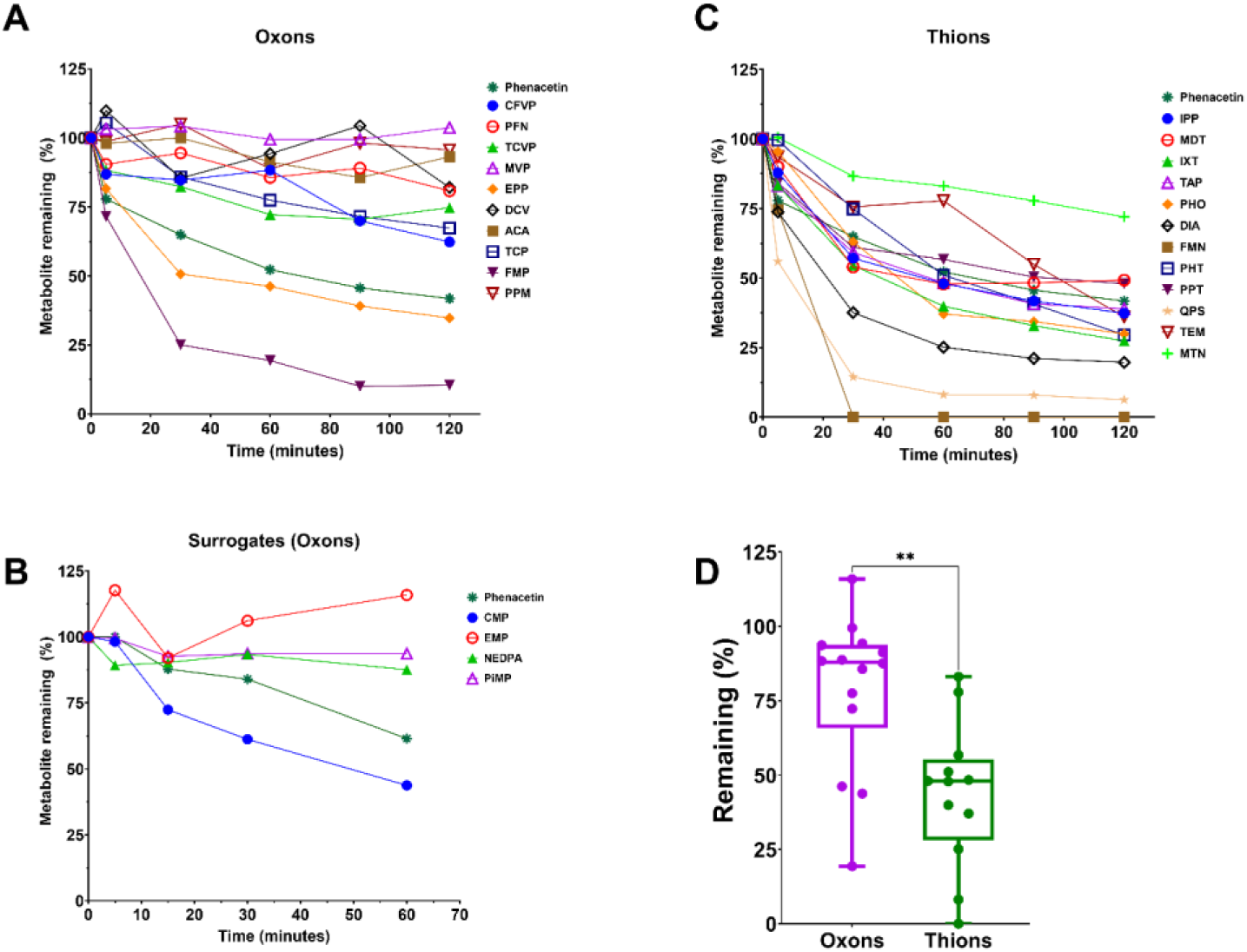
Metabolism of OP compounds by CYP1A2 measured by disappearance of the parent. **A**. LC-MS/MS quantification of the degradation of oxons by CYP1A2 after 2 hrs. **B**. LC-MS/MS quantification of the degradation of thions by CYP1A2 after 2 hrs. **C**. LC-MS/MS quantification of the degradation of CWNA surrogates after 1 hr. **D. CYP1A2-mediated degradation of thions was significantly higher than that of oxons based on mean values after 1 hr.** **P<0.01. Dark green dots represent thions; violet dots represent oxons.

**Table 1.** Half-life and intrinsic clearance of OP compounds by CYP1A2 as measured by disappearance of parent.

| Compound | t<sub>1/2</sub> (min) | CL<sub>int</sub> (μL/min/pmol CYP) |
| --- | --- | --- |
| Phenacetin | 104.3 | 0.1 |
| **Oxon** |  |  |
| PFN, DCV, ACA, MVP, PPM | >372.8 | <0.04 |
| PiMP, EMP, NEDPA | >186.4 | <0.07 |
| CFVP | 210.8 | 0.1 |
| TCVP | 194.8 | 0.1 |
| TCP | 190.4 | 0.1 |
| EPP | 32.3 | 0.4 |
| CMP | 49.4 | 0.28 |
| FMP | 28.8 | 0.5 |
| **Thion** |  |  |
| IPP | 87.5 | 0.2 |
| MDT | 55.0 | 0.3 |
| IXT | 66.3 | 0.2 |
| TAP | 91.1 | 0.2 |
| PHO | 65.3 | 0.2 |
| DIA | 41.1 | 0.3 |
| FMN | 11.9 | 1.2 |
| PHT | 66.9 | 0.2 |
| PPT | 45.5 | 0.3 |
| TEM | 91.4 | 0.2 |
| QPS | 17.1 | 0.8 |
| MTN | 254.4 | 0.1 |

### Molecular docking analysis shows correlations between the individual docking parameters and fluorogenic assay-based inhibition or LC-MS/MS-based degradation experiments

Rigid docking in the CYP1A2 receptor (PDB ID: 2HI4) was performed using AutoDock Vina with each of the oxons or thions (Supplementary Figure S1 and S2). 2HI4 has one of the best resolutions among the CYP1A2 crystal structures and has been used extensively for docking previously (Sansen *et al*., 2007; Zhu *et al*., 2011; Watanabe *et al*., 2017; Dutkiewicz and Mikstacka, 2018; Navapour and Mogharrab, 2021) as it contains α-naphthoflavone bound to the heme center. We continued the correlation analysis between the experimental results of the fluorogenic assay-based inhibition and LC-MS/MS metabolism experiments and the *in silico* docking parameters for OP compounds, as was done previously with CYP2C9 (Shriwas, Noonchester, Revnew, *et al*., 2026) and CYP3A4 (Shriwas, Noonchester, Scarpitti, *et al*., 2026). Here, we also included correlations with various molecular parameters for each OP compound, such as molecular weight (MW), among others (Supplementary Table 1), just as in the case of CYP3A4, along with the docking parameters (binding energy (BE), reactive distance (D), and binding efficiency (BEeff)) for the best pose and the pose with the shortest reactive distance (SD, BE(SD), BEeff(SD)) (Supplementary Table 2). Correlation was determined between each of the individual molecular and docking parameters (Supplementary Figure S3) and significance levels (Supplementary Table 11). Interestingly, we found that the experimental data from the LC-MS/MS metabolism and pFluor50 fluorogenic assay inhibition analyses were correlated with each other. We found that the % OP compound remaining by LC-MS/MS (2 hours) had a negative correlation (R = -0.71; P<0.001) with the 10 µM 30-minute inhibition data and a positive correlation with time-dependent inhibition at 10 µM (R = 0.51; P<0.05) (Figure 3A-B). Furthermore, the % OP remaining by LC-MS/MS (1 hour) had a negative correlation (R = -0.60; P<0.01) with the 10 µM 30-minute inhibition data and a negative correlation with the 1 µM 10- minute inhibition data (R = -0.54; P<0.01) (Figure 3C-D). Additionally, the 10 µM 10-minute inhibition had a positive correlation with the molecular weight (R = 0.56; P<0.0001), while the time-dependent inhibition at 10 µM had a positive correlation (R = 0.36; P<0.05) with BEeff (SD) (Figure 3E-F). Similarly, other significant correlations between experimental parameters such as pFluor50-based inhibition and LC-MS/MS metabolism are reported (Supplementary Figure S4) in which the % OP compound remaining after 2 hours correlated with the time-dependent inhibition at 1 µM (R = -0.50; P<0.05) and with the 1 µM 10-minute inhibition (R = - 0.50; P<0.05), and correlated negatively with the 10 µM 10-minute inhibition (R = -0.47; P<0.05) (Supplementary Figure S4A-C). Further, the % OP compound remaining after 1 hour of metabolism was significantly negatively correlated with the 10 µM 10-minute inhibition (R = - 0.53; P<0.01) and time-dependent inhibition at 1 µM (R = -0.46; P<0.05) (Supplementary Figure S4D-E). We also report significant correlations between docking parameters and experimental data (Supplementary Figure S5). Interestingly, BEeff (SD) was correlated with the 10 µM 10- minute inhibition (R = 0.32; P<0.05), 1 µM 10-minute inhibition (R = 0.32, P<0.05), and time-dependent inhibition with at 1 µM (R = 0.30; P<0.05) (Supplementary Figure S5A,D-E). Further, the 10 µM 10-minute inhibition with BEeff (D) (R = 0.35; P<0.05) and the 1 µM 30-minute inhibition with D (R = -0.32; P<0.05) were also significantly correlated (Supplementary Figure S5B-C). Overall, all correlations with their R and P values are reported (Supplementary Tables 4–10).

**Figure 3.**
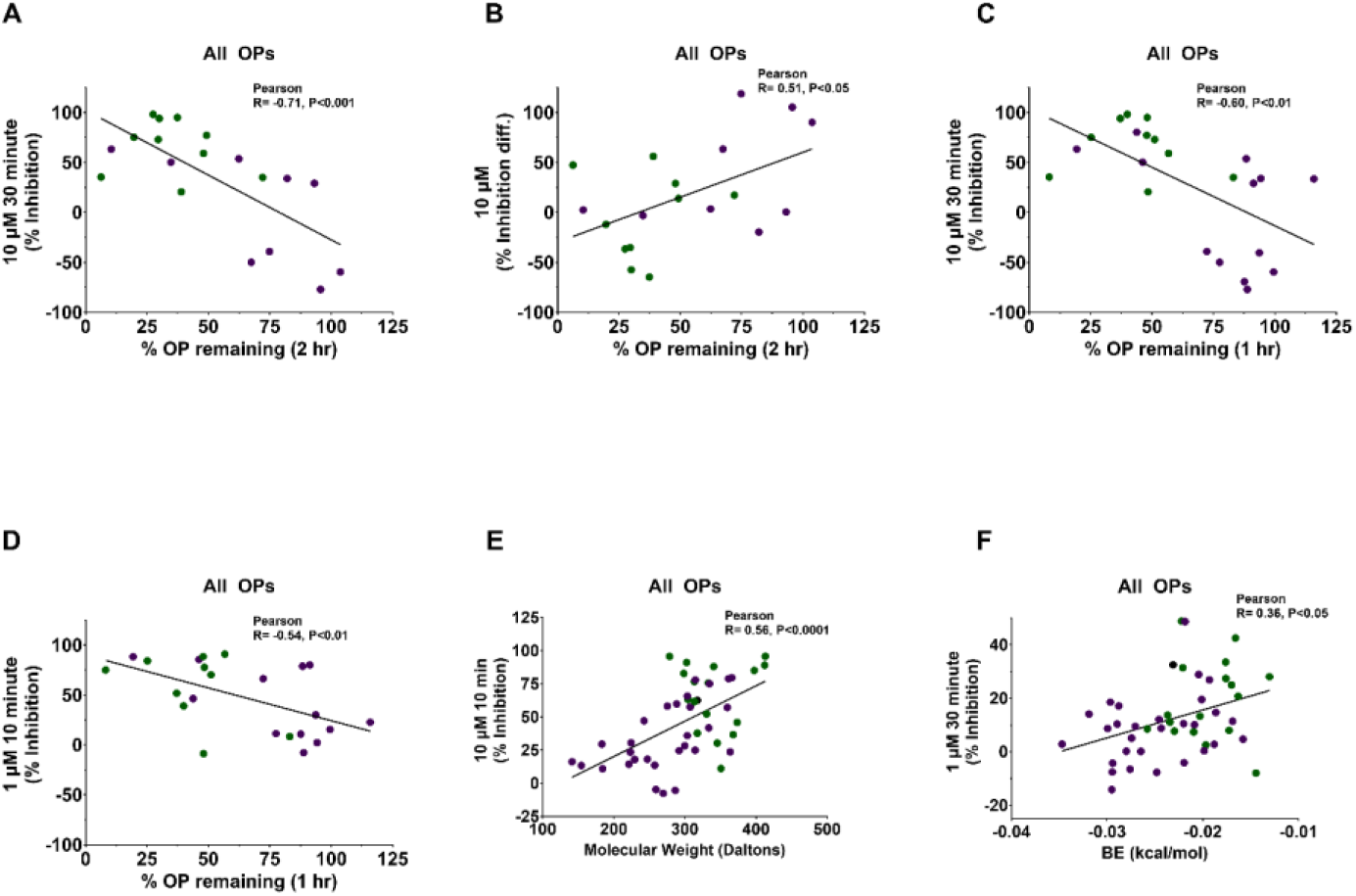
**Correlation analysis between molecular parameters, pFluor50-based CYP1A2 inhibition data, and LC-MS/MS data. A-F**. Correlation analysis between inhibition at 10 µM after 30-minute preincubation with % OP compound remaining (2 hours) (Pearson R = -0.71; P<0.001), % OP compound remaining (2 hours) and time-dependent inhibition at 10 µM (Pearson R = 0.51; P<0.05), inhibition at 10 µM after 30-minute preincubation with % OP compound remaining (1 hour) (Pearson R = -0.60; P<0.01), % OP compound remaining (1 hour) and inhibition at 1 µM after 10-minute preincubation (Pearson R = -0.54; P<0.01), inhibition at 10 µM after 10-minute preincubation with molecular weight (Pearson R = 0.56; P<0.0001), time- dependent inhibition at 10 µM with Binding energy effective (Pearson R = 0.36; P<0.05). Dark green dots represent thions; violet dots represent oxons.

We also explored the interactions between CYP1A2 and the CWNA surrogates compared with the authentic CWNAs when docked into the 2HI4 structure. Many residues in 2HI4 formed common interactions with both the CWNAs and the surrogates (Supplementary Figure S6 A-H). For some surrogate–CWNA pairs, such as CMP and cyclosarin, and NEDPA and tabun (Supplementary Figure S6 A-B, G-H), many residues in the CYP1A2 receptor were involved in common interactions. However, the other pairs, PiMP and soman as well as EMP and VX, did not share similar interacting residues (Supplementary Figure S6C-F). This again suggests that CYP1A2 may discriminate between the surrogates and their corresponding CWNAs, and that metabolism of the authentic CWNAs needs to be validated experimentally rather than inferred from these docking studies. Further, the 3D docked poses (shortest distance) showed that cyclosarin and its surrogate CMP interacted similarly, whereas sarin differed in both the R and S configurations (Figure 4). Overall, because CMP was degraded by CYP1A2 and interacts with the enzyme similarly to cyclosarin, it is possible that CYP1A2 also degrades cyclosarin.

**Figure 4.**
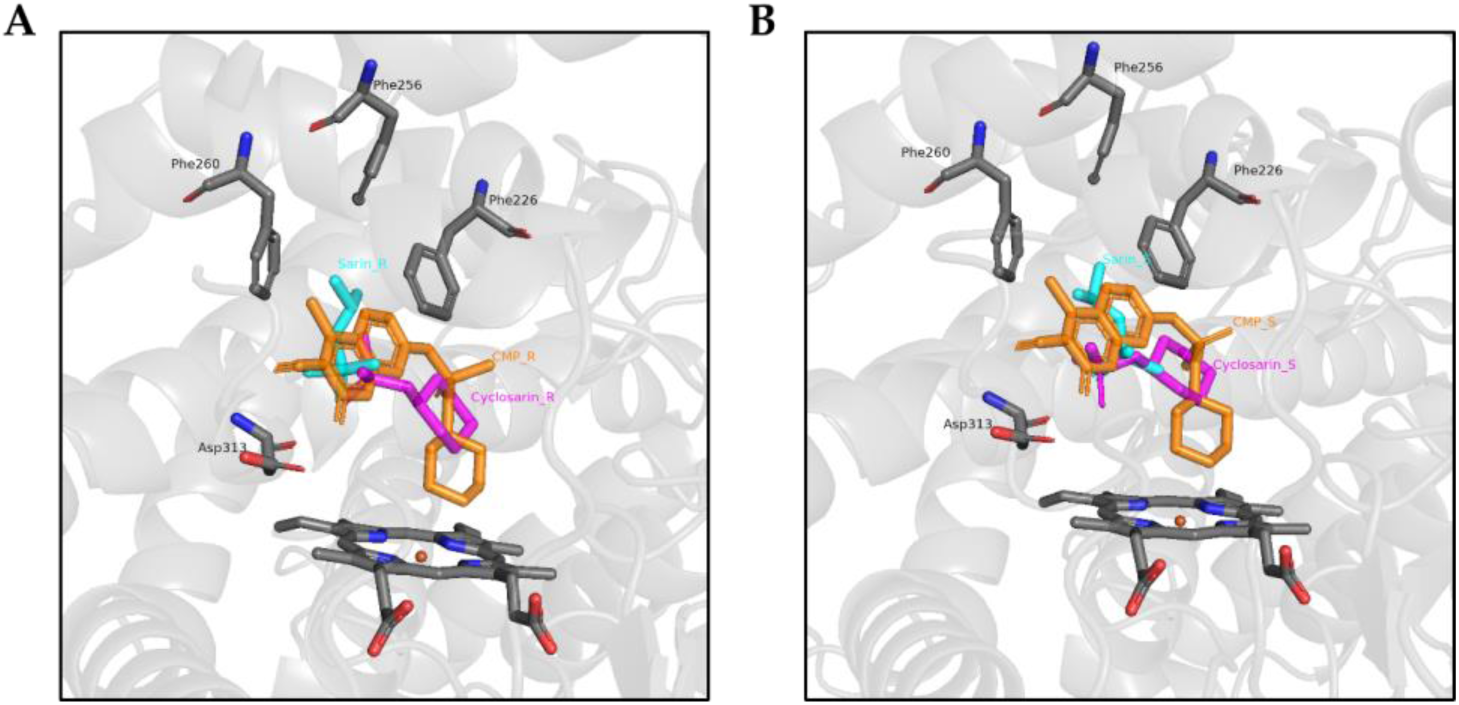
3D docking poses (shortest reactive distance) for the R and S enantiomers of CMP/cyclosarin/sarin with 2HI4 PDB structure of CYP1A2 **A.** 3D docking poses for CMP (*R*), cyclosarin (*R*) and sarin (*R*)**. B.** 3D docking poses for CMP (*S*), cyclosarin (*S*) and sarin (*S*).

## Discussion

OP poisoning has become a global health concern, partly owing to the high levels of exposure among agricultural laborers as well as to pesticide residues in treated crops and vegetables. The fatality rate is 10–20% per incident (Eddleston *et al*., 2008), nearly 44% of exposed farmers report poisoning symptoms (Damalas and Koutroubas, 2016) and approximately 41% of food samples analyzed in the European Union in 2022 contained quantifiable pesticide residues ((European Food Safety Authority (EFSA) *et al*., 2024), which could lead to toxicity. The lack of effective therapies against these toxic agents also contributes to the high mortality of intentional OP self-poisoning, and there is therefore an urgent need for medical countermeasures against OP poisoning (Lovins *et al*., 2024, 2026).

We previously developed the pFluor50 fluorogenic assay for quantifying CYP1A2 activity using Eres as a fluorogenic substrate, which was validated using α-naphthoflavone as an inhibitor of CYP1A2 activity (Shriwas *et al*., 2025). In this study, we used this assay to test activation and inhibition by OP compounds in a time- (10 and 30 minutes) and dose-dependent (1 and 10 µM) manner. Many OP compounds such as CFVP, EPP, FMP, IBF, MCP and NEDPA were inhibitors at 1 µM after 10 minutes but became activators after 30 minutes. Many OP compounds at 10 µM, such as CPO, CFA, and CTP, among others, showed increased inhibition at 30 minutes, whereas their inhibition at 1 µM decreased between 10 and 30 minutes (Figure 1A-B). These variations in activity point to possible mechanism-based inhibition or stronger binding to the CYP1A2 enzyme. Interestingly, time-dependent inhibition differences were seen for thions (more variations at 10 µM) and oxons (more variations at 1 µM). This further highlights the selectivity of CYP1A2 towards thions at higher concentrations. Overall, these results align with previous reports showing CYP1A2 to be a metabolizer of thions such as CPY, DIA, APM, and PE (Buratti *et al*., 2003; Ellison *et al*., 2012) and with reports that genetic polymorphisms in CYP1A2 may modify susceptibility to pesticide toxicity (Chilipweli *et al*., 2025).

The time-dependent changes in pFluor50 activity can be interpreted in more than one way. Increased inhibition between 10 and 30 minutes, as seen for CPO, CFA, and CTP at 10 µM, is consistent with mechanism-based inhibition, degradation to a more inhibitory metabolite, or slow, tight binding. The decrease in inhibition at 1 µM, and the apparent activation after 30 minutes seen with CFVP, EPP, FMP, IBF, MCP, and NEDPA, could instead reflect depletion of the parent OP by CYP1A2 during preincubation, or the formation of metabolites that inhibit less than the parent compound. Because only two preincubation times were tested, IC50-shift or kinetic/KI experiments, together with metabolite identification, will be needed to distinguish these mechanisms.

Our previous NMR-based work, and that of others on OP compounds, has shown that many are metabolized in a single-phase or biphasic manner by HLMs (Sams, Mason and Rawbone, 2000; Buratti *et al*., 2005; Rose *et al*., 2005; Ellison *et al*., 2012; Agarwal *et al*., 2023). Because of the large number of enzymes present in HLMs, it is difficult to determine which ones are the true metabolizers of these OP compounds. We therefore previously examined the individual enzymes CYP2C9, CYP2C19, and CYP3A4 (Vignaux *et al*., 2023; Shriwas, Noonchester, Revnew, *et al*., 2026; Shriwas, Noonchester, Scarpitti, *et al*., 2026). However, many OP compounds were not metabolized by any of these three, so in this study we investigated whether CYP1A2 could directly degrade OP compounds using LC-MS/MS. Further, CYP1A2 is highly selective, metabolizing only ∼10% of drugs, making it interesting to investigate whether it is similarly selective towards OP compounds. Thions were more likely to be degraded than oxons, with a significantly lower % remaining after 1 hour of metabolism (Figure 2D). Consistent with previous reports, DIA was metabolized by CYP1A2 in our study, which supports the validity of our approach (Ellison *et al*., 2012) (Figure 2B).

The kinetic parameters derived from the parent-disappearance experiments place the observed metabolism in quantitative context (Table 1). Phenacetin, the canonical CYP1A2 probe substrate, was cleared with a t1/2 of 104.3 minutes and a CLint of 0.1 µL/min/pmol CYP under our conditions. Several OP compounds were cleared substantially faster than this positive control, including the thions FMN (11.9 min; 1.2) and QPS (17.1 min; 0.8), and the oxons FMP (28.8 min; 0.5) and EPP (32.3 min; 0.4). At the other extreme, the oxons PFN, DCV, ACA, MVP and PPM showed no measurable loss of parent over 2 hours (t1/2 > 372.8 min; CLint < 0.04), and MTN was the slowest metabolized thion (254.4 min; 0.1). That the best OP substrates exceed the clearance of a diagnostic CYP1A2 substrate supports regarding this isoform, which constitutes 13–15% of hepatic CYP450 content, as a potentially meaningful contributor to OP compound disposition rather than an incidental one, although intrinsic clearance by a recombinant enzyme does not by itself predict hepatic clearance. The spread of more than an order of magnitude in CLint across a structurally diverse compound set also indicates that the determinants of turnover are structural rather than simply a function of the P=S or P=O substitution.

An important caveat applies to the interpretation of preferential thion metabolism. Our LC-MS/MS assay measures loss of the parent compound, which for a P=S compound reports the sum of oxidative desulfuration and dearylation. The former yields the corresponding oxon, a more potent covalent inhibitor of AChE than the parent thion, whereas the latter is a genuine detoxification route (Bharate et al., 2010; Thompson, Prins and George, 2010). CYP1A2 has been reported to catalyze both transformations for OP pesticides such as CPY and DIA (Tang et al., 2001; Ellison et al., 2012), so the rapid disappearance of thions observed here cannot by itself be equated with detoxification, and the balance between the two pathways will determine whether CYP1A2 acts as a bioscavenger or a bioactivator for any given substrate. The comparatively slow clearance of most oxons is relevant in this regard, since an oxon generated *in situ* would not necessarily be efficiently removed by the same enzyme. Resolving this will require identification and quantification of the individual metabolites and direct measurement of their AChE inhibitory potency, which we identify below as the principal next step.

We then performed docking analysis with the 2HI4 receptor structure and found that many docking parameters correlated with the experimental results from pFluor50 inhibition and LC-MS/MS metabolomics. Notably, the 10 µM 30-minute inhibition showed a significant negative correlation with the % OP compound remaining by LC-MS/MS at 2 hours (R = -0.71; P<0.001) and 1 hour (R = -0.60, P<0.01). Other correlations were also observed and are tabulated in Supplementary Tables 9 and 10. We also found that docking parameters were correlated with the experimental data. In fact, BEeff (SD) showed a significant correlation with the time-dependent inhibition at 10 µM (R = 0.36; P<0.05), 10 µM 10-minute inhibition (R = 0.32; P<0.05), 1 µM 10-minute inhibition (R = 0.32; P<0.05) and time-dependent inhibition at 1 µM (R = 0.30; P<0.05) (Figure 3F; Supplementary Figure S5 A, D-E). We have tabulated all correlations between docking and molecular properties for all time- and dose-dependent inhibition data (Supplementary Table 3-10).

Several features of the docking and correlation analyses should be interpreted cautiously. The positive correlation between molecular weight and 10-minute inhibition at 10 µM (R = 0.56; P<0.0001) suggests that larger OP compounds inhibit CYP1A2 more strongly, whereas the weaker correlations of BEeff at the shortest reactive distance with the inhibition measures (R = 0.30–0.36) indicate that rigid-receptor docking captures only part of what determines inhibition.

CYP1A2 has a narrow, planar active site that favors small, planar, aromatic ligands (Sansen *et al*., 2007) which may help explain why several aromatic thions, such as QPS, PPT, and DIA, were cleared rapidly, but we did not test this directly. In addition, many parameter pairs were correlated without correction for multiple comparisons, so the correlations are best regarded as hypothesis-generating.

The inverse correlation between inhibition at 10 µM (30 minutes) and the percentage of OP compound remaining is mechanistically plausible: a compound that is turned over by CYP1A2 must bind in its active site and can therefore compete with the probe substrate Eres. If confirmed with additional compounds, this relationship suggests that the high-throughput pFluor50 assay could be used to prioritize OP comopunds for the more resource-intensive LC-MS/MS clearance assay. The correlation is moderate, however, and inhibition can also arise from tight binding without turnover, as with several oxons that inhibited CYP1A2 but were not degraded, so the fluorogenic assay cannot replace direct measurement of metabolism.

Finally, we investigated the 2D interactions of the CWNA surrogates and the authentic CWNAs (Figure S6). CMP and the CWNA cyclosarin had similar interacting residues such as Phe226, Asp313, and Asn312, although there were some differences (Ser122, Thr118), and 3D docking also placed them at similar binding sites with Phe226, Phe256, Phe260, and Asp313 (Figure S6A-B; Figure 4). These results point to the need to investigate the direct metabolism of cyclosarin by CYP1A2 and are consistent with the selectivity of CYP1A2 towards CWNAs.

Our study is limited in that it tested only direct OP compound degradation; the identity of the secondary metabolites produced remains to be investigated, and those metabolites would need to be tested for AChE toxicity. In future work, we will also explore more rigorous models such as Molecular Mechanics Poisson–Boltzmann Surface Area (MM-PBSA), which may provide additional parameters for correlation and for developing an AI/ML model of the metabolism of these and other OP compounds by CYP1A2. In addition, all experiments used a single recombinant enzyme preparation *in vitro* at one (LC-MS/MS) or two (pFluor50) fixed OP compound concentrations and only two incubation times (10 and 30 minutes), so enzyme kinetic parameters (Km and Vmax) were not determined and extrapolation to hepatic clearance *in vivo* remains untested. Route of exposure also matters: hepatic CYP1A2 may contribute to the clearance of OP compounds after ingestion or systemic absorption, but whether it could act quickly enough to protect against inhaled CWNAs is unknown. Known interindividual variability in CYP1A2 activity, such as genetic polymorphisms and induction by tobacco smoke, and competition from other OP compounds and other metabolic routes in the liver were also not examined. Finally, docking used a rigid receptor and a single crystal structure, which may not capture the conformational flexibility of the CYP1A2 active site.

In conclusion, recombinant human CYP1A2 degraded a substantial subset of the tested OP compounds, most efficiently thions, and several OP compounds inhibited its activity in a time- and concentration-dependent manner. Inhibition and metabolism data were correlated and docking placed cyclosarin and its surrogate CMP in similar binding poses, supporting direct testing of cyclosarin metabolism. Whether CYP1A2 acts as a bioscavenger or a bioactivator for individual OP compounds will depend on the metabolites formed and their AChE inhibitory potency. At present, these findings identify CYP1A2 as a candidate contributor to OP compound metabolism rather than an established bioscavenger. Together, these data provide a foundation for AI/ML models that predict CYP1A2-mediated metabolism of existing and emerging OP compounds.

## Patents

Not applicable.

## Supplementary Materials

Supplementary materials are attached separately.

## Author Contributions

Conceptualization, P.S. and C.M.; methodology, P.S. and A.N.; software, P.S., B.S., A.N., and C.H.; validation, C.M., C.H., T.L. and S.E.; formal analysis, P.S. and C.M.; investigation, P.S., B.S., A.R., and A.N.; resources, C.M., C.H. and S.E.; data curation, P.S.; writing—original draft preparation, P.S.; writing—review and editing, P.S., C.M., C.H., T.L. and S.E.; visualization, P.S.; supervision, C.M., C.H., and S.E.; project administration, C.M. and S.E.; funding acquisition, C.M. and S.E. All authors have read and agreed to the published version of the manuscript.

## Institutional Review Board Statement

Not applicable.

## Informed Consent Statement

Not applicable.

## Data Availability Statement

Data are contained within the article and the Supplementary Materials.

## Acknowledgments

The authors would like to thank Dmitry Uchenik (Shared Instrumentation Facility Director, College of Pharmacy) for use of the BioTek plate reader. Computational resources were provided by the Ohio Supercomputer Center.

## Abbreviations

AChE, Acetylcholinesterase; ACA, Acephate; APM, Azinphos-methyl; BE, binding energy; BNS, Bensulide; CFVP, Chlorfenvinphos; CMP, cyclosarin surrogate; CPH, Chlorphoxim; CPY, Chlorpyrifos; CPO, Chlorpyrifos Oxon; CTP, Crotoxyphos; CFA, Crufomate; CFP, Cyanofenphos; CYP, cytochrome P450; CWNA, chemical warfare nerve agent; DFP, Diisopropyl fluorophosphate; DIA, Diazinon; DCV, Dichlorvos;; DMX, Dimefox; DMA, Dimethoate; DMP, Dimethyl paraoxon; EMP, VX surrogate; EPP, Ethoprophos; Eres, resorufin ethoxy ether; FMP, Fenamiphos; FNN, Fenthion; FON, Formothion; IFP, Iodofenphos; IBF, Iprobenfos; IPP, Isofenphos; IXT, Isoxathion; LPS, Leptophos; MTN, Malathion; MPF, Mephosfolan; MMP, Methamidophos; MDT, Methidathion; MVP, Mevinphos; MCP, Monocrotophos; NEDPA, Tabun surrogate; OP, organophosphorus compound; EP, Diethylparaoxon; PHO, Phosalone; PiMP, Soman surrogate; PFN, Phosfolan; PHT, Phosmet; PPM, Phosphamidon; PPE, Pirimiphos-ethyl; PZP, Pyrazophos; PPT, Pyridaphenthion; QPS, Quinalphos; RRB, Rotatable bonds; SCN, Schradan; TEM, Temephos; TCVP, Tetrachlorvinphos; TAP, Triazophos; TBS, Tribufos; TCP, Trichlorphon.

