## Supplementary for "Human CYP1A2 metabolism of organophosphorus pesticides and nerve agent surrogates"

### Supplementary Material

#### Supplementary Figures

**Supplementary Figure 1. Structures of the oxons used in experiments and docking analysis**

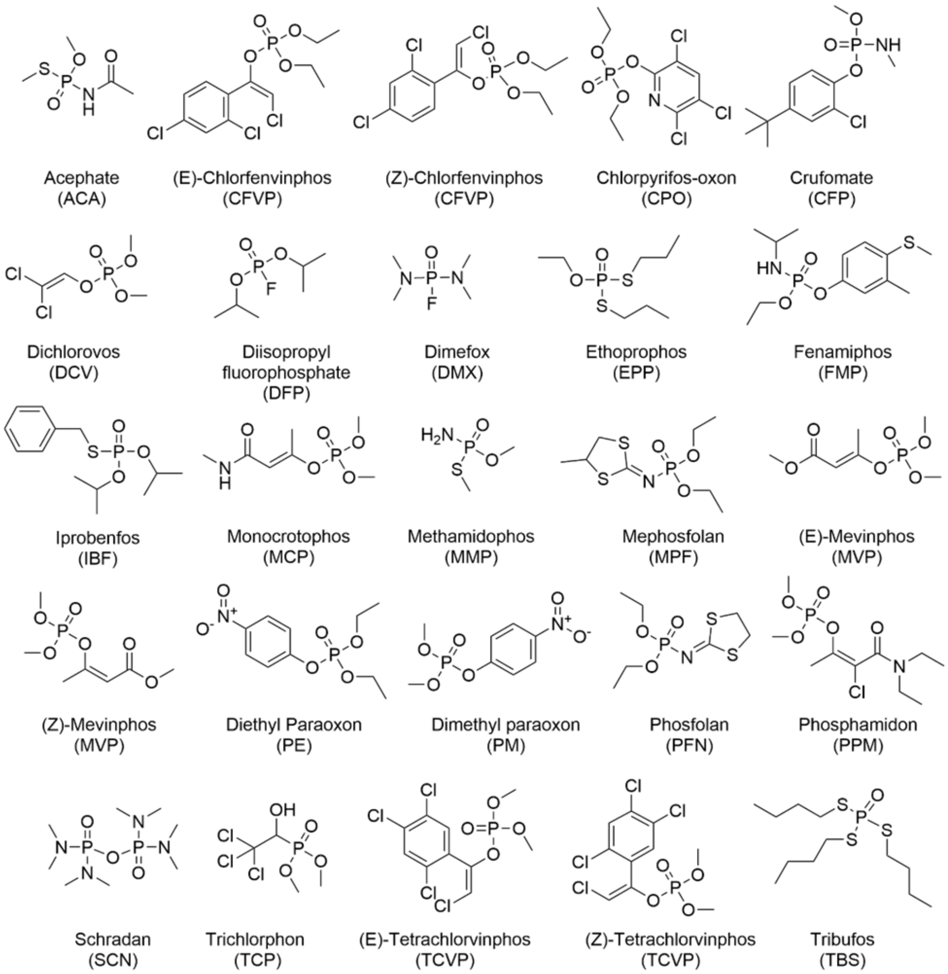


Structures of the oxons used in experiments and docking analysis

C2 - Confidential Business Data

**Supplementary Figure 2. Structures of the thions used in experiments and docking analysis**

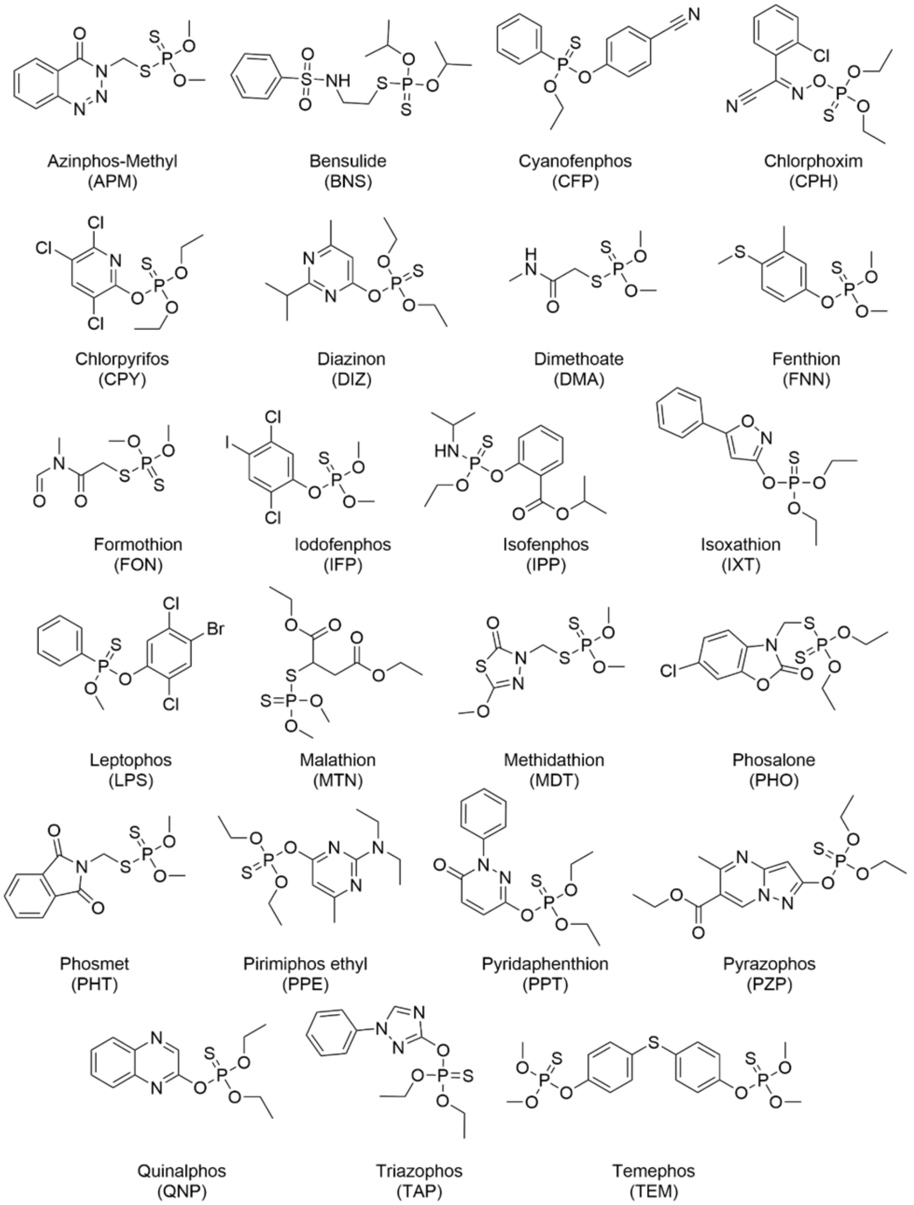


Structures of the thions used in experiments and docking analysis

**Supplementary Figure S3. Correlation values between molecular and docking parameters for CYP1A2 (2HI4 pdb)**

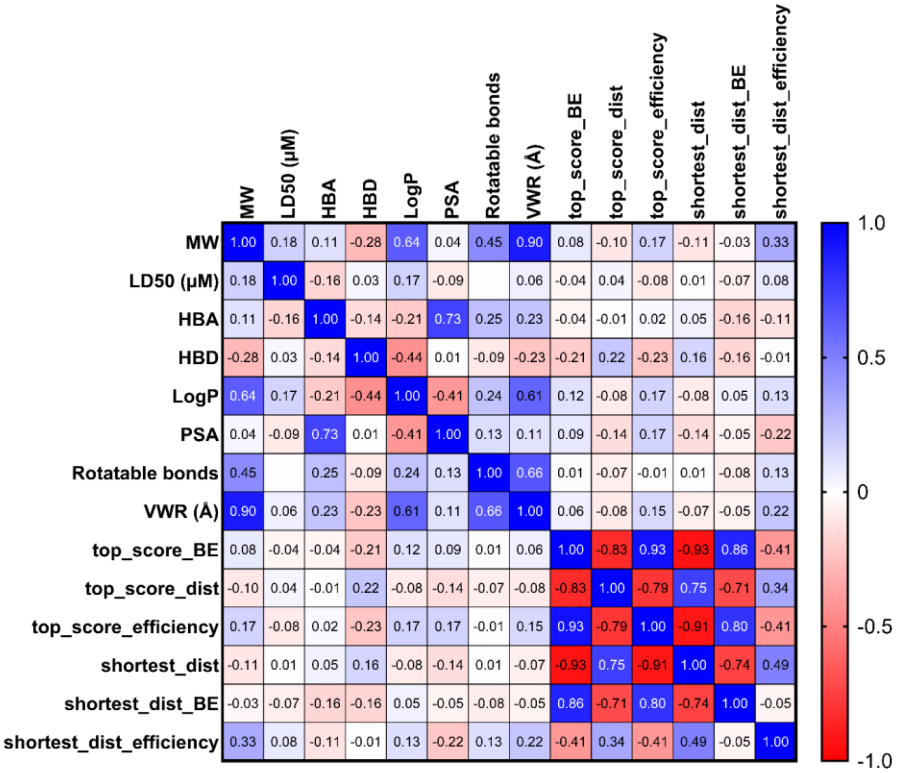


**Supplementary Figure S3. Correlation values between molecular and docking parameters for CYP1A2.** Correlation matrix between docking parameters and molecular parameters for the 47 OP compounds used in the paper. Averages were taken for R/S stereocenters or E/Z geometric isomeric pairs for OP compounds with stereocenters/geometric isomers. The blue color indicates strong positive correlation while the red color indicates strong negative correlation.

C2 - Confidential Business Data

**Supplementary Figure S4. Correlation analysis between LC-MS/MS metabolism at 1 and 2 hours with pFlour50 mediated inhibition**

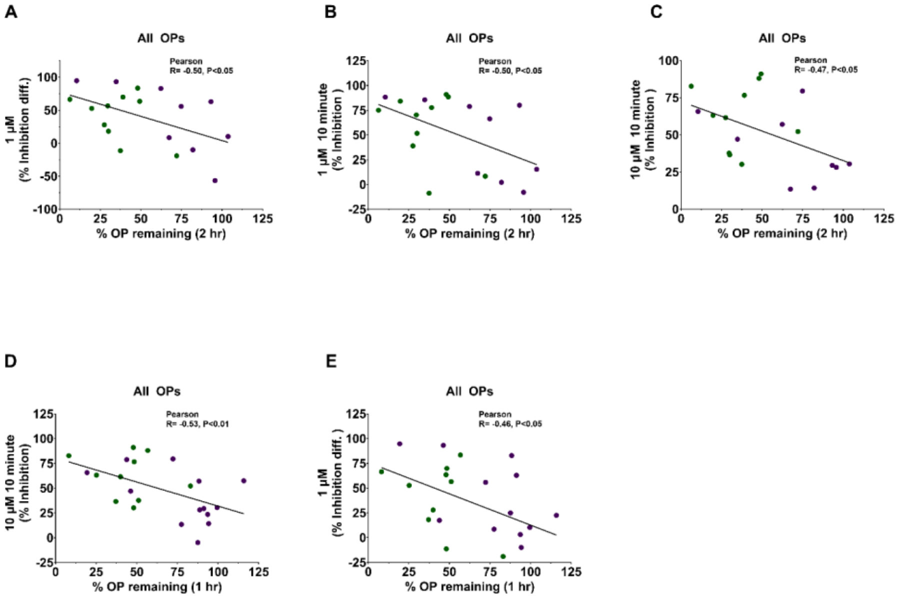


**Supplementary Figure S4. Pearson Correlation analysis between pFlour50 mediated CYP1A2 inhibition with % OP remaining after 1 or 2 hours (LC-MS/MS metabolism). A.** Correlation analysis (R = 0.50, P<0.05) between % of OP remaining after 2 hours and time-dependent inhibition for all OP compounds at 1 µM. **B**. Correlation analysis (R = 0.50, P<0.05) between % of OP compound remaining after 2 hours and inhibition by all OP compounds at 1 µM with10 min preincubation. **C**. Correlation analysis (R = –0.47, P<0.05) between % of OP compound remaining after 2 hours and inhibition by all OP compounds at 10 µM with 10 min preincubation. **D**. Correlation analysis (R = –0.53, P<0.01) between % of OP compound remaining after 1 hour and inhibition by all OP compounds at 10 µM with 10 min preincubation. **E**. Correlation analysis (R = –0.46, P<0.05) between % of OP compound remaining after 1 hour and time-dependent inhibition for all OP compounds at 1 µM.

C2 - Confidential Business Data

**Supplementary Figure S5. Correlation analysis between docking parameters with pFlour50 mediated inhibition**

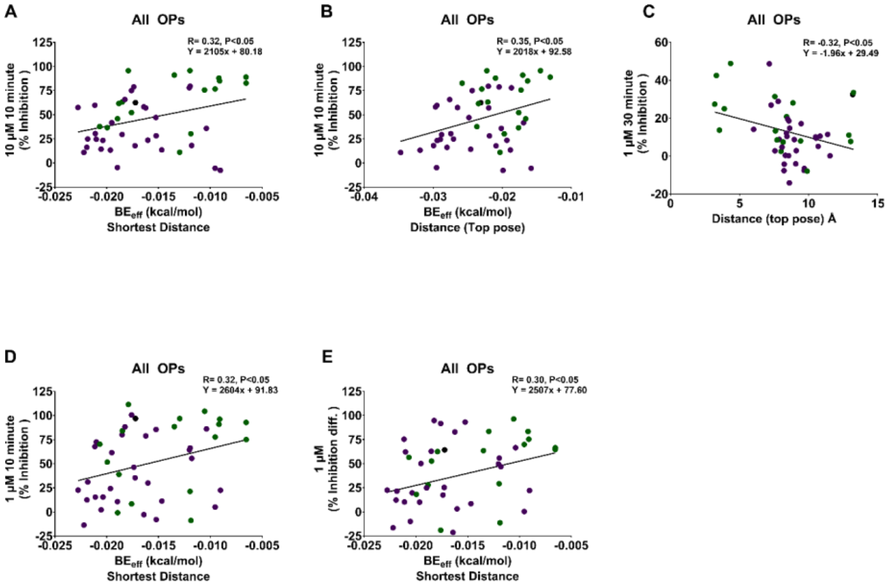


**Supplementary Figure S5. Pearson Correlation analysis between pFlour50 mediated CYP1A2 inhibition with docking parameters. A.** Correlation analysis (R = 0.33, P<0.05) between $BE_{eff}$ (SD) and inhibition by all OP compounds at 10 µM with 10 min preincubation. **B**. Correlation analysis (R = 0.33, P<0.05) between $BE_{eff}$ (D) and inhibition by all OP compounds at 10 µM with 10 min preincubation. **C**. Correlation analysis (R = –0.31, P<0.05) between Distance (top pose) and inhibition by all OP compounds at 1 µM with 30 min preincubation. **D**. Correlation analysis (R = 0.32, P<0.05) between $BE_{eff}$ (SD) and inhibition by all OP compounds at 1 µM with 10 min preincubation. **E**. Correlation analysis (R = 0.30, P<0.05) between $BE_{eff}$ (SD) and time-dependent inhibition by all OP compounds at 1 µM.

**Supplementary Figure S6. 2D interacting partners between CWNA surrogates and CWNA with 2HI4 PDB structure of CYP1A2**

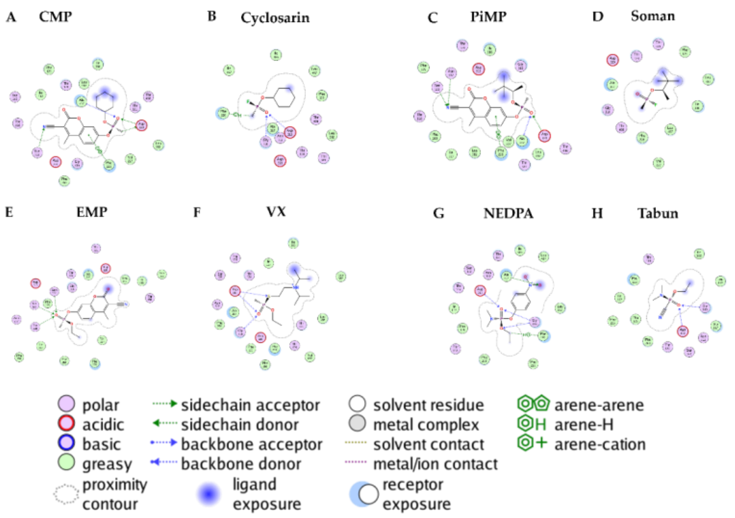


**Supplementary Figure S6. 2D interaction diagrams in R conformation for CWNA and CWNA surrogates. A**. CMP (R) interacted with groups such as Phe226, Asp320 and Asn312 among others. **B**. 2D interaction diagram of Cyclosarin (R) with Phe226, Gly326, and Asp313 among others being closely interacting residues. **C**. 2D interaction diagram of PiMP (R) with Phe226, Asp320 and Asn312 among other closely interacting residues. **D**. 2D interaction diagram of Soman (R) with Ala317, Gly326, and Asp320 among other interacting residues. **E**. EMP (R) interacts with Thr118, Asn312 and Ser122 among others. **F**. 2D interaction diagram of VX (R) with Asp313 and Gly316 among other interacting residues. **G**. 2D interaction diagram of NEDPA (R) with Asp313 and Gly316 among other interacting residues. **H**. 2D interaction diagram of Tabun (R) with Asp313 and Gly316 or GlyB316 among other interacting residues. Legends panel for various interactions is provided.

#### Supplementary Tables

**Supplementary Table 1: Table of molecular properties of the OP compounds**

| OP Compound | MW | $LD_{50}$ (µM) | HBA | HBD | LogP | PSA | Rotatable bonds | VWR (Å) |
|---|---|---|---|---|---|---|---|---|
| ACA | 183.17 | 163.88[32, 33] | 2 | 1 | -0.0006 | 55.4 | 4 | 153 |
| APM | 317.32 | 0.90[32, 33] | 3 | 0 | 2.72 | 63.49 | 5 | 261.625 |
| BNS | 397.51 | 21.58[32, 33] | 2 | 1 | 0 | 64.63 | 10 | 352.5 |
| CFVP | 359.57 | 1.06[32, 33] | 1 | 0 | 3.4575 | 44.76 | 7 | 272.125 |
| CPH | 332.74 | 238.66[32, 33] | 2 | 0 | 4.6882 | 63.84 | 8 | 281.75 |
| CPY | 350.59 | 5.98[32, 33] | 1 | 0 | 4.9668 | 40.05 | 6 | 305.9375 |
| CPO | 334.50 | 6.27[32, 33] | 2 | 0 | 4.2326 | 57.12 | 6 | 328.375 |
| CMP | 361.33 | 0.32[38, 39] | 3 | 0 | 3.016 | 85.62 | 5 | 291 |
| CTP | 314.27 | -0.2 | 2 | 0 | 2.3893 | 71.06 | 8 | 269.75 |
| CFA | 291.71 | 103.45[32, 33] | 1 | 1 | 3.532 | 47.56 | 5 | 282.75 |
| CFP | 303.30 | 4.50[32, 33] | 1 | 0 | 4.4521 | 42.25 | 6 | 292.625 |
| DIZ | 304.35 | 3.32[32, 33] | 2 | 0 | 5.0438 | 52.41 | 7 | 170.625 |
| DCV | 220.98 | 11.50[32, 33] | 1 | 0 | 1.8444 | 44.76 | 4 | 234.25 |
| DEP | 275.19 | 0.35[32, 33] | 2 | 0 | 1.697 | 96.57 | 7 | 165.5 |
| DFP | 184.15 | 1.04[32, 33] | 1 | 0 | 2.2165 | 35.53 | 4 | 286.375 |
| DMX | 154.13 | 0.21[32, 33] | 1 | 0 | 0.5649 | 23.55 | 2 | 192 |
| DMA | 229.26 | 33.95[32, 33] | 1 | 1 | 0.445 | 47.56 | 6 | 198.5 |
| DMP | 247.14 | 0.26[32, 33] | 2 | 0 | 0.975 | 96.57 | 5 | 139 |
| EMP | 307.24 | 0.38[34, 35] | 3 | 0 | 1.7734 | 85.62 | 5 | 269.875 |
| EPP | 242.34 | 5.24[32, 33] | 1 | 0 | 3.2058 | 26.3 | 8 | 229.75 |
| FMP | 303.36 | 0.63[32, 33] | 1 | 1 | 3.401 | 47.56 | 7 | 292.75 |
| FNN | 278.33 | 3.64[32, 33] | 0 | 0 | 4.09 | 27.69 | 5 | 247.125 |
| FON | 257.30 | 30.86[32, 33] | 2 | 0 | 0.4522 | 55.84 | 7 | 280.5 |
| IFP | 413.00 | 179.21[32, 33] | 0 | 0 | 5.16 | 27.69 | 4 | 263 |
| IBF | 288.34 | 74.91[32, 33] | 1 | 0 | 4.1856 | 35.53 | 7 | 328.375 |
| IPP | 345.39 | 2.58[32, 33] | 1 | 1 | 4.053 | 56.79 | 9 | 279.625 |
| IXT | 313.31 | 11.36[32, 33] | 2 | 0 | 4.3074 | 49.28 | 7 | 286.125 |
| LPS | 412.06 | 3.32[32, 33] | 0 | 0 | 5.9015 | 18.46 | 4 | 197.625 |
| MTN | 330.36 | 170.96[32, 33] | 2 | 0 | 1.8403 | 71.06 | 11 | 233.875 |
| MPF | 269.30 | 31.85[32, 33] | 1 | 0 | 2.9563 | 47.89 | 5 | 114.375 |

**Commented [CM1]:** Each of these properties should have the same number of decimal places or the same number of significant figures amongst all the entries… not sure why it is changing. My guess is that it is an artifact of Excel which doesn't know what significant figures are so it erases zeros, but please fix.

**Formatted:** Centered, Space Before: 3 pt, After: 3 pt, Line spacing: single

**Formatted Table**

**Formatted:** Space Before: 3 pt, After: 3 pt, Line spacing: single

**Formatted** ...

**Formatted** ...

**Formatted** ...

**Formatted** ...

**Formatted** ...

**Formatted** ...

**Formatted** ...

**Formatted** ...

**Formatted** ...

**Formatted** ...

**Formatted** ...

**Formatted** ...

**Formatted** ...

**Formatted** ...

**Formatted** ...

**Formatted** ...

**Formatted** ...

**Formatted** ...

**Formatted** ...

**Formatted** ...

**Formatted** ...

**Formatted** ...

**Formatted** ...

**Formatted** ...

**Formatted** ...

**Formatted** ...

**Formatted** ...

**Formatted** ...

**Formatted** ...

C2 - Confidential Business Data

| OP Compound | MW | $LD_{50}$ (µM) | HBA | HBD | LogP | PSA | Rotatable bonds | VWR (Å) |
|---|---|---|---|---|---|---|---|---|
| MMP | 141.13 | 6.75[32, 33] | 1 | 1 | -0.66 | 52.32 | 2 | 232.75 |
| MDT | 302.33 | 2.63[32, 33] | 3 | 0 | 3.2045 | 60.36 | 6 | 286.625 |
| MVP | 224.15 | 0.50[32, 33] | 2 | 0 | 0.1996 | 71.06 | 6 | 194.0625 |
| MCP | 223.20 | 1.99[32, 33] | 2 | 1 | 999.4683 | 73.86 | 6 | 245.75 |
| NEDPA | 259.092 | 27.52[36, 37] | 2 | 0 | 1.699 | 90.58 | 6 | 307.25 |
| PHO | 367.81 | 10.36[32, 33] | 1 | 0 | 4.4271 | 48 | 7 | 260.125 |
| PFN | 255.30 | 1.11[32, 33] | 1 | 0 | 2.5432 | 47.89 | 5 | 342.75 |
| PHT | 317.32 | 11.31[32, 33] | 2 | 0 | 2.805 | 55.84 | 5 | 317.75 |
| PPM | 299.69 | 0.74[32, 33] | 2 | 0 | 0.3829 | 65.07 | 8 | 251.875 |
| PiMP | 363.50 | 0.97[34, 38] | 3 | 0 | 3.5634 | 85.62 | 6 | 304.625 |
| PPE | 333.39 | 13.34[32, 33] | 3 | 0 | 4.4999 | 55.65 | 9 | 327.25 |
| PZP | 373.36 | 12.85[32, 33] | 4 | 0 | 3.4558 | 81.95 | 9 | 270 |
| PPT | 340.33 | 71.78[32, 33] | 2 | 0 | 1.836 | 60.36 | 7 | 266.875 |
| QNP | 298.30 | 7.56[32, 33] | 2 | 0 | 3.5306 | 52.41 | 6 | 279.25 |
| SCN | 286.25 | 0.56[32, 33] | 2 | 0 | 0.716 | 56.33 | 6 | 315.375 |
| TEM | 466.50 | 68.09[32, 33] | 0 | 0 | 5.6665 | 55.38 | 10 | 192.5 |
| TCVP | 365.95 | 347.20[32, 33] | 1 | 0 | 4.8974 | 44.76 | 5 | 290.625 |
| TAP | 313.31 | 6.69[32, 33] | 3 | 0 | 3.781 | 55.65 | 7 | 153 |
| TBS | 314.50 | 3.12[32, 33] | 1 | 0 | 5.2066 | 17.07 | 12 | 261.625 |
| TCP | 257.44 | 26.16[32, 33] | 2 | 1 | 0.473333 | 55.76 | 4 | 352.5 |

**Formatted:** Centered, Space Before: 3 pt, After: 3 pt, Line spacing: single

**Formatted Table**

**Formatted:** Space Before: 3 pt, After: 3 pt, Line spacing: single

**Formatted:** Space Before: 3 pt, After: 3 pt, Line spacing: single

**Formatted:** Space Before: 3 pt, After: 3 pt, Line spacing: single

**Formatted:** Space Before: 3 pt, After: 3 pt, Line spacing: single

**Formatted:** Space Before: 3 pt, After: 3 pt, Line spacing: single

**Formatted:** Space Before: 3 pt, After: 3 pt, Line spacing: single

**Formatted:** Space Before: 3 pt, After: 3 pt, Line spacing: single

**Formatted:** Space Before: 3 pt, After: 3 pt, Line spacing: single

**Formatted:** Space Before: 3 pt, After: 3 pt, Line spacing: single

**Formatted:** Space Before: 3 pt, After: 3 pt, Line spacing: single

**Formatted:** Space Before: 3 pt, After: 3 pt, Line spacing: single

**Formatted:** Space Before: 3 pt, After: 3 pt, Line spacing: single

**Formatted:** Space Before: 3 pt, After: 3 pt, Line spacing: single

**Formatted:** Space Before: 3 pt, After: 3 pt, Line spacing: single

**Formatted:** Space Before: 3 pt, After: 3 pt, Line spacing: single

**Formatted:** Space Before: 3 pt, After: 3 pt, Line spacing: single

**Formatted:** Space Before: 3 pt, After: 3 pt, Line spacing: single

**Formatted:** Space Before: 3 pt, After: 3 pt, Line spacing: single

**Formatted:** Space Before: 3 pt, After: 3 pt, Line spacing: single

**Formatted:** Space Before: 3 pt, After: 3 pt, Line spacing: single

C2 - Confidential Business Data

**Supplementary Table 2: Table of docking properties of the OP compounds**

| OP Compound | BE (D) | D | $BE_{eff}$ (D) | (SD) | BE(SD) | $BE_{eff}$(SD) |
|---|---|---|---|---|---|---|
| ACA | -4.980 | 9.458 | -0.029 | 4.230 | -3.205 | -0.019 |
| APM | -7.049 | 13.188 | -0.023 | 2.946 | -5.255 | -0.017 |
| BNS | -6.089 | 8.405 | -0.016 | 3.237 | -3.392 | -0.009 |
| CFA | -7.888 | 8.426 | -0.029 | 3.372 | -5.725 | -0.021 |
| CFP | -5.818 | 8.479 | -0.020 | 5.172 | -2.999 | -0.010 |
| CFVP | -7.580 | 9.065 | -0.022 | 5.602 | -5.608 | -0.016 |
| CMP | -6.972 | 7.795 | -0.020 | 7.588 | -5.931 | -0.017 |
| CPH | -5.501 | 9.419 | -0.017 | 3.496 | -3.361 | -0.011 |
| CPO | -7.869 | 7.876 | -0.024 | 6.440 | -5.693 | -0.018 |
| CPY | -6.896 | 7.595 | -0.020 | 4.839 | -4.396 | -0.013 |
| CTP | -7.980 | 10.534 | -0.027 | 3.161 | -6.442 | -0.022 |
| DCV | -5.253 | 8.387 | -0.025 | 3.809 | -4.394 | -0.021 |
| DEP | -5.901 | 9.097 | -0.035 | 4.331 | -3.775 | -0.022 |
| DFP | -6.251 | 7.553 | -0.022 | 4.685 | -5.237 | -0.018 |
| DIZ | -4.535 | 10.505 | -0.021 | 3.166 | -3.748 | -0.017 |
| DMA | -4.530 | 6.018 | -0.032 | 4.130 | -2.790 | -0.020 |
| DMP | -7.755 | 11.544 | -0.026 | 2.894 | -6.674 | -0.023 |
| DMX | -5.535 | 8.216 | -0.025 | 4.284 | -3.402 | -0.015 |
| EMP | -7.758 | 9.683 | -0.028 | 4.563 | -5.129 | -0.018 |
| EPP | -5.847 | 4.350 | -0.022 | 3.224 | -4.713 | -0.018 |
| FMP | -7.869 | 9.671 | -0.029 | 4.757 | -5.649 | -0.021 |
| FNN | -5.847 | 9.892 | -0.014 | 3.426 | -4.842 | -0.012 |
| IBF | -6.324 | 7.998 | -0.020 | 5.144 | -3.813 | -0.012 |
| IFP | -6.963 | 12.909 | -0.023 | 3.186 | -5.594 | -0.019 |
| IPP | -5.243 | 8.861 | -0.013 | 6.180 | -2.631 | -0.007 |
| IXT | -5.478 | 3.198 | -0.018 | 3.198 | -5.478 | -0.018 |
| LPS | -6.148 | 8.229 | -0.029 | 3.208 | -4.256 | -0.020 |
| MCP | -4.954 | 3.895 | -0.017 | 3.207 | -3.924 | -0.013 |
| MDT | -3.723 | 8.548 | -0.028 | 3.184 | -2.915 | -0.022 |
| MMP | -5.790 | 10.688 | -0.027 | 3.868 | -4.452 | -0.021 |
| MPF | -5.031 | 8.313 | -0.020 | 5.977 | -2.285 | -0.009 |

**Formatted:** Space Before: 3 pt, After: 3 pt

C2 - Confidential Business Data

| | | | | | | |
|---|---|---|---|---|---|---|
| MTN | -7.503 | 8.591 | -0.027 | 4.262 | -5.430 | -0.020 |
| MVP | -7.742 | 8.564 | -0.030 | 6.201 | -3.958 | -0.016 |
| NEDPA | -7.642 | 8.602 | -0.030 | 5.057 | -4.913 | -0.019 |
| PHO | -7.222 | 3.542 | -0.024 | 3.819 | -4.901 | -0.021 |
| PHT | -6.589 | 7.280 | -0.019 | 4.663 | -4.878 | -0.016 |
| PiMP | -7.097 | 8.956 | -0.030 | 3.442 | -3.644 | -0.012 |
| PPE | -5.221 | 11.376 | -0.017 | 4.219 | -5.474 | -0.020 |
| PPM | -6.128 | 7.153 | -0.022 | 3.383 | -4.920 | -0.015 |
| PPT | -6.775 | 8.150 | -0.021 | 2.971 | -3.234 | -0.009 |
| PZP | -5.860 | 3.321 | -0.017 | 7.997 | -6.464 | -0.019 |
| QNP | -7.304 | 7.708 | -0.026 | 6.848 | -1.710 | -0.007 |
| SCN | -4.140 | 8.074 | -0.016 | 6.211 | -2.737 | -0.010 |
| TAP | -6.821 | 13.036 | -0.023 | 6.211 | -2.737 | -0.010 |
| TBS | -5.357 | 8.622 | -0.019 | 3.661 | -2.997 | -0.012 |
| TCP | -4.691 | 7.557 | -0.019 | 3.207 | -5.238 | -0.015 |
| TCVP | -7.848 | 10.824 | -0.022 | 3.721 | -4.503 | -0.012 |

**Formatted:** Space Before: 3 pt, After: 3 pt

**Formatted:** Space Before: 3 pt, After: 3 pt
**Formatted:** Space Before: 3 pt, After: 3 pt
**Formatted:** Space Before: 3 pt, After: 3 pt
**Formatted:** Space Before: 3 pt, After: 3 pt
**Formatted:** Space Before: 3 pt, After: 3 pt
**Formatted:** Space Before: 3 pt, After: 3 pt
**Formatted:** Space Before: 3 pt, After: 3 pt
**Formatted:** Space Before: 3 pt, After: 3 pt
**Formatted:** Space Before: 3 pt, After: 3 pt
**Formatted:** Space Before: 3 pt, After: 3 pt
**Formatted:** Space Before: 3 pt, After: 3 pt
**Formatted:** Space Before: 3 pt, After: 3 pt
**Formatted:** Space Before: 3 pt, After: 3 pt
**Formatted:** Space Before: 3 pt, After: 3 pt
**Formatted:** Space Before: 3 pt, After: 3 pt

**Supplementary Table 3. Correlation between the time-dependent ~~difference in~~ inhibition at 1 µM (10 ~~and~~ or 30 minutes preincubation) with the molecular parameters**

| Property | R | P (two-tailed) | Slope | Y-intercept |
|---|---|---|---|---|
| MW | 0.212 | 0.153 | 0.346 | 286.100 |
| LD50 (µM) | 0.162 | 0.277 | 0.302 | 23.720 |
| HBA | -0.191 | 0.199 | -0.004 | 1.868 |
| HBD | 0.016 | 0.914 | 1.69E-04 | 0.185 |

**Formatted:** Font: Bold

**Formatted:** Space Before: 3 pt, After: 3 pt, Line spacing: single

**Formatted:** Space Before: 3 pt, After: 3 pt, Line spacing: single

**Formatted:** Space Before: 3 pt, After: 3 pt, Line spacing: single

**Formatted:** Space Before: 3 pt, After: 3 pt, Line spacing: single

**Formatted:** Space Before: 3 pt, After: 3 pt, Line spacing: single

C2 - Confidential Business Data

| | | | | |
|---|---|---|---|---|
| LogP | -0.065 | 0.665 | -0.247 | 33.330 |
| *PSA | -0.348 | 0.016 | -0.178 | 62.220 |
| Rotatable bonds | 0.015 | 0.918 | 0.001 | 6.225 |
| VWR (Å) | 0.217 | 0.143 | 0.312 | 247.500 |
| BE (D) | -0.270 | 0.066 | -0.008 | -5.950 |
| D | 0.195 | 0.188 | 0.012 | 8.135 |
| $BE_{eff}$ (D) | 0.043 | 0.774 | 5.67E-06 | -0.023 |
| SD | -0.050 | 0.738 | -0.002 | 4.413 |
| BE(SD) | 0.175 | 0.239 | 0.006 | -4.610 |
| *$BE_{eff}$(SD) | 0.300 | 0.041 | 3.60E-05 | -0.017 |

Parameters with * are significantly correlated with experimental result

Formatted: Space Before: 3 pt, After: 3 pt, Line spacing: single

Formatted: Space Before: 3 pt, After: 3 pt, Line spacing: single

Formatted: Space Before: 3 pt, After: 3 pt, Line spacing: single

Formatted: Space Before: 3 pt, After: 3 pt, Line spacing: single

Formatted: Space Before: 3 pt, After: 3 pt, Line spacing: single

Formatted: Space Before: 3 pt, After: 3 pt, Line spacing: single

Formatted: Space Before: 3 pt, After: 3 pt, Line spacing: single

Formatted: Space Before: 3 pt, After: 3 pt, Line spacing: single

Formatted: Space Before: 3 pt, After: 3 pt, Line spacing: single

Formatted: Space Before: 3 pt, After: 3 pt, Line spacing: single

C2 - Confidential Business Data

**Supplementary Table 4. Correlation between the time-dependent ~~difference in~~ inhibition at 10 µM (10 ~~and~~ or 30 minutes preincubation) with molecular parameters**

| Property | R | P (two-tailed) | Slope | Y-intercept |
|---|---|---|---|---|
| MW | -0.091 | 0.544 | -0.128 | 301.100 |
| LD50 (µM) | 0.138 | 0.353 | 0.223 | 31.740 |
| HBA | 0.334 | 0.022 | 0.007 | 1.602 |
| HBD | -0.049 | 0.746 | 4.36E-04 | 0.198 |
| LogP | 0.224 | 0.131 | 0.733 | 13.040 |
| PSA | 0.461 | 0.001 | 0.204 | 52.460 |
| Rotatable bonds | -0.057 | 0.703 | -0.003 | 6.295 |
| VWR (Å) | -0.131 | 0.382 | -0.162 | 261.700 |
| BE | 0.023 | 0.879 | 0.001 | -6.261 |
| D | -0.144 | 0.333 | -0.008 | 8.708 |
| $BE_{eff}$ | -0.037 | 0.804 | 4.49E-03 | -0.023 |
| SD | 0.150 | 0.316 | 0.004 | 4.280 |
| BE(SD) | 0.125 | 0.403 | 0.004 | -4.449 |
| $BE_{eff}$(SD) | 0.176 | 0.236 | 1.82E-05 | -0.016 |

Parameters with * are significantly correlated with experimental result

**Formatted:** Font: Bold

**Formatted:** Space Before: 3 pt, After: 3 pt, Line spacing: single

**Formatted:** Space Before: 3 pt, After: 3 pt, Line spacing: single

**Formatted:** Space Before: 3 pt, After: 3 pt, Line spacing: single

**Formatted:** Space Before: 3 pt, After: 3 pt, Line spacing: single

**Formatted:** Space Before: 3 pt, After: 3 pt, Line spacing: single

**Formatted:** Space Before: 3 pt, After: 3 pt, Line spacing: single

**Formatted:** Space Before: 3 pt, After: 3 pt, Line spacing: single

**Formatted:** Space Before: 3 pt, After: 3 pt, Line spacing: single

**Formatted:** Space Before: 3 pt, After: 3 pt, Line spacing: single

**Formatted:** Space Before: 3 pt, After: 3 pt, Line spacing: single

**Formatted:** Space Before: 3 pt, After: 3 pt, Line spacing: single

**Formatted:** Space Before: 3 pt, After: 3 pt, Line spacing: single

**Formatted:** Space Before: 3 pt, After: 3 pt, Line spacing: single

**Formatted:** Space Before: 3 pt, After: 3 pt, Line spacing: single

**Formatted:** Space Before: 3 pt, After: 3 pt, Line spacing: single

C2 - Confidential Business Data

**Supplementary Table 5. Correlation between the inhibition at 1 µM ~~at~~ with 10 minute~~s~~ preincubation with the molecular parameters**

| **Property** | **R** | **P (two-tailed)** | **Slope** | **Y-intercept** |
|---|---|---|---|---|
| *MW | 0.320 | 0.028 | 0.536 | 272.200 |
| LD50 (µM) | 0.123 | 0.410 | 0.236 | 23.240 |
| HBA | -0.104 | 0.487 | -0.002 | 1.826 |
| HBD | -0.073 | 0.627 | -0.001 | 0.231 |
| LogP | -0.134 | 0.370 | -0.521 | 50.250 |
| *PSA | -0.325 | 0.026 | -0.171 | 64.100 |
| Rotatable bonds | 0.076 | 0.610 | 0.004 | 6.045 |
| *VWR (Å) | 0.316 | 0.030 | 0.467 | 235.800 |
| BE | -0.227 | 0.124 | -0.007 | -5.903 |
| D | 0.076 | 0.612 | 0.005 | 8.348 |
| $BE_{eff}$ | 0.181 | 0.224 | 2.44E-05 | -0.024 |
| SD | -0.026 | 0.860 | -0.001 | 4.395 |
| BE(SD) | 0.123 | 0.408 | 0.004 | -4.603 |
| *$BE_{eff}$(SD) | 0.324 | 0.027 | 3.98E-05 | -0.018 |

Parameters with * are significantly correlated with experimental result; Pearson Correlation

**Formatted:** Font: Bold

**Formatted:** Space Before: 3 pt, After: 3 pt, Line spacing: single

**Formatted:** Space Before: 3 pt, After: 3 pt, Line spacing: single

**Formatted:** Space Before: 3 pt, After: 3 pt, Line spacing: single

**Formatted:** Space Before: 3 pt, After: 3 pt, Line spacing: single

**Formatted:** Space Before: 3 pt, After: 3 pt, Line spacing: single

**Formatted:** Space Before: 3 pt, After: 3 pt, Line spacing: single

**Formatted:** Space Before: 3 pt, After: 3 pt, Line spacing: single

**Formatted:** Space Before: 3 pt, After: 3 pt, Line spacing: single

**Formatted:** Space Before: 3 pt, After: 3 pt, Line spacing: single

**Formatted:** Space Before: 3 pt, After: 3 pt, Line spacing: single

**Formatted:** Space Before: 3 pt, After: 3 pt, Line spacing: single

**Formatted:** Space Before: 3 pt, After: 3 pt, Line spacing: single

**Formatted:** Space Before: 3 pt, After: 3 pt, Line spacing: single

**Formatted:** Space Before: 3 pt, After: 3 pt, Line spacing: single

C2 - Confidential Business Data

**Supplementary Table 6. Correlation between the inhibition at 1 µM ~~at~~ with 30 minute~~s~~ preincubation with the molecular parameters**

| Property | R | P (two-tailed) | Slope | Y-intercept |
|---|---|---|---|---|
| MW | 0.262 | 0.075 | 1.121 | 284.900 |
| LD50 (µM) | -0.109 | 0.464 | -0.534 | 41.870 |
| HBA | 0.234 | 0.114 | 0.014 | 1.523 |
| HBD | -0.228 | 0.123 | -0.006 | 0.270 |
| LogP | -0.171 | 0.249 | -1.703 | 45.670 |
| PSA | 0.081 | 0.588 | 0.109 | 54.140 |
| Rotatable bonds | 0.155 | 0.299 | 0.022 | 5.982 |
| VWR (Å) | 0.240 | 0.104 | 0.904 | 247.800 |
| BE | 0.126 | 0.399 | 0.010 | -6.377 |
| *D | -0.317 | 0.030 | -0.052 | 9.246 |
| *$BE_{eff}$ | 0.349 | 0.016 | 1.20E-04 | -0.024 |
| SD | 0.064 | 0.671 | 0.006 | 4.274 |
| BE(SD) | -0.143 | 0.337 | -0.012 | -4.242 |
| $BE_{eff}$(SD) | 0.041 | 0.782 | 1.30E-05 | -0.016 |

Parameters with * are significantly correlated with experimental result; Pearson Correlation

**Formatted:** Font: Bold

**Formatted:** Space Before: 3 pt, After: 3 pt, Line spacing: single

**Formatted:** Space Before: 3 pt, After: 3 pt, Line spacing: single

**Formatted:** Space Before: 3 pt, After: 3 pt, Line spacing: single

**Formatted:** Space Before: 3 pt, After: 3 pt, Line spacing: single

**Formatted:** Space Before: 3 pt, After: 3 pt, Line spacing: single

**Formatted:** Space Before: 3 pt, After: 3 pt, Line spacing: single

**Formatted:** Space Before: 3 pt, After: 3 pt, Line spacing: single

**Formatted:** Space Before: 3 pt, After: 3 pt, Line spacing: single

**Formatted:** Space Before: 3 pt, After: 3 pt, Line spacing: single

**Formatted:** Space Before: 3 pt, After: 3 pt, Line spacing: single

**Formatted:** Space Before: 3 pt, After: 3 pt, Line spacing: single

**Formatted:** Space Before: 3 pt, After: 3 pt, Line spacing: single

**Formatted:** Space Before: 3 pt, After: 3 pt, Line spacing: single

**Formatted:** Space Before: 3 pt, After: 3 pt, Line spacing: single

**Formatted:** Space Before: 3 pt, After: 3 pt, Line spacing: single

**Supplementary Table 7. Correlation between inhibition at 10 µM with 30 minutes preincubation with molecular parameters**

| Property | R | P (two-tailed) | Slope | Y-intercept |
|---|---|---|---|---|
| *MW | 0.370 | 0.010 | 0.418 | 286.000 |
| LD50 (µM) | 0.015 | 0.923 | 0.019 | 34.500 |
| HBA | -0.257 | 0.081 | -0.004 | 1.831 |
| HBD | -0.072 | 0.630 | -0.001 | 0.208 |
| LogP | -0.238 | 0.107 | -0.625 | 43.710 |
| *PSA | -0.464 | 0.001 | -0.164 | 60.680 |
| Rotatable bonds | 0.183 | 0.219 | 0.007 | 6.044 |
| *VWR (Å) | 0.355 | 0.014 | 0.353 | 248.200 |
| BE | -0.162 | 0.278 | -0.003 | -6.148 |
| D | 0.123 | 0.409 | 0.005 | 8.425 |
| $BE_{eff}$ | 0.208 | 0.160 | 1.89E-05 | -0.023 |
| SD | -0.088 | 0.555 | -0.002 | 4.414 |
| BE(SD) | -0.114 | 0.447 | -0.003 | -4.316 |
| $BE_{eff}$(SD) | 0.037 | 0.803 | 3.10E-06 | -0.016 |

Parameters with * are significantly correlated with experimental result; Spearman Correlation

Formatted: Line spacing: single

Formatted: Font: Bold

Formatted: Space Before: 3 pt, After: 3 pt, Line spacing: single

Formatted: Space Before: 3 pt, After: 3 pt, Line spacing: single

Formatted: Space Before: 3 pt, After: 3 pt, Line spacing: single

Formatted: Space Before: 3 pt, After: 3 pt, Line spacing: single

Formatted: Space Before: 3 pt, After: 3 pt, Line spacing: single

Formatted: Space Before: 3 pt, After: 3 pt, Line spacing: single

Formatted: Space Before: 3 pt, After: 3 pt, Line spacing: single

Formatted: Space Before: 3 pt, After: 3 pt, Line spacing: single

Formatted: Space Before: 3 pt, After: 3 pt, Line spacing: single

Formatted: Space Before: 3 pt, After: 3 pt, Line spacing: single

Formatted: Space Before: 3 pt, After: 3 pt, Line spacing: single

Formatted: Space Before: 3 pt, After: 3 pt, Line spacing: single

Formatted: Space Before: 3 pt, After: 3 pt, Line spacing: single

Formatted: Space Before: 3 pt, After: 3 pt, Line spacing: single

**Supplementary Table 8. Correlation between inhibition at 10 µM 10 minutes with molecular parameters**

| **Property** | **R** | **P (two-tailed)** | **Slope** | **Y-intercept** |
|---|---|---|---|---|
| *MW | 0.555 | <0.0001 | 1.169 | 244.800 |
| LD50 (µM) | 0.234 | 0.114 | 0.562 | 8.969 |
| HBA | 0.019 | 0.897 | 0.001 | 1.675 |
| HBD | -0.207 | 0.163 | -0.003 | 0.320 |
| LogP | -0.110 | 0.460 | -0.540 | 49.160 |
| PSA | -0.178 | 0.232 | -0.117 | 60.960 |
| Rotatable bonds | 0.256 | 0.083 | 0.018 | 5.440 |
| *VWR (Å) | 0.467 | 0.001 | 0.866 | 219.000 |
| BE | -0.268 | 0.069 | -0.010 | -5.776 |
| D | 0.015 | 0.921 | 0.001 | 8.536 |
| *$BE_{eff}$ | 0.333 | 0.022 | 5.4E-05 | -0.025 |
| SD | 0.058 | 0.698 | 0.003 | 4.226 |
| BE(SD) | -0.025 | 0.866 | -0.001 | -4.347 |
| *$BE_{eff}$(SD) | 0.333 | 0.022 | 5.14E-05 | -0.018 |

Parameters with * are significantly correlated with experimental result; Spearman Correlation

Formatted: Line spacing: single

Formatted: Font: Bold

Formatted: Space Before: 3 pt, After: 3 pt, Line spacing: single

Formatted: Space Before: 3 pt, After: 3 pt, Line spacing: single

Formatted: Space Before: 3 pt, After: 3 pt, Line spacing: single

Formatted: Space Before: 3 pt, After: 3 pt, Line spacing: single

Formatted: Space Before: 3 pt, After: 3 pt, Line spacing: single

Formatted: Space Before: 3 pt, After: 3 pt, Line spacing: single

Formatted: Space Before: 3 pt, After: 3 pt, Line spacing: single

Formatted: Space Before: 3 pt, After: 3 pt, Line spacing: single

Formatted: Space Before: 3 pt, After: 3 pt, Line spacing: single

Formatted: Space Before: 3 pt, After: 3 pt, Line spacing: single

Formatted: Space Before: 3 pt, After: 3 pt, Line spacing: single

Formatted: Space Before: 3 pt, After: 3 pt, Line spacing: single

Formatted: Space Before: 3 pt, After: 3 pt, Line spacing: single

Formatted: Space Before: 3 pt, After: 3 pt, Line spacing: single

Formatted: Line spacing: single

**Supplementary Table 9. Correlation between LC-MS/MS 2 hours degradation with molecular parameters**

| **Property** | **R** | **P (two-tailed)** | **Slope** | **Y-intercept** |
|---|---|---|---|---|
| MW | -0.332 | 0.131 | -0.676 | 336.900 |
| LD50 (µM) | 0.308 | 0.163 | 0.851 | 0.905 |
| HBA | 0.008 | 0.973 | 1.82E-04 | 1.627 |
| HBD | 0.033 | 0.883 | 4.34E-05 | 0.160 |
| *LogP | -0.512 | 0.015 | -0.029 | 4.227 |
| PSA | 0.302 | 0.172 | 0.097 | 48.660 |
| Rotatable bonds | -0.262 | 0.239 | -0.016 | 7.475 |
| *VWR (Å) | -0.442 | 0.039 | -0.826 | 302.100 |
| BE | 0.168 | 0.454 | 0.006 | -6.396 |
| D | -0.065 | 0.774 | -0.006 | 8.783 |
| $BE_{eff}$ | -0.207 | 0.356 | -3.10E-05 | -0.020 |
| SD | -0.265 | 0.234 | -0.010 | 4.496 |
| BE(SD) | -0.150 | 0.505 | -0.005 | -4.076 |
| $BE_{eff}$(SD) | -0.186 | 0.407 | -2.48E-05 | -0.014 |

Parameters with * are significantly correlated with experimental result; Pearson Correlation

**Formatted:** Font: Bold
**Formatted:** Space Before: 3 pt, After: 3 pt, Line spacing: single
**Formatted:** Space Before: 3 pt, After: 3 pt, Line spacing: single
**Formatted:** Space Before: 3 pt, After: 3 pt, Line spacing: single
**Formatted:** Space Before: 3 pt, After: 3 pt, Line spacing: single
**Formatted:** Space Before: 3 pt, After: 3 pt, Line spacing: single
**Formatted:** Space Before: 3 pt, After: 3 pt, Line spacing: single
**Formatted:** Space Before: 3 pt, After: 3 pt, Line spacing: single
**Formatted:** Space Before: 3 pt, After: 3 pt, Line spacing: single
**Formatted:** Space Before: 3 pt, After: 3 pt, Line spacing: single
**Formatted:** Space Before: 3 pt, After: 3 pt, Line spacing: single
**Formatted:** Space Before: 3 pt, After: 3 pt, Line spacing: single
**Formatted:** Space Before: 3 pt, After: 3 pt, Line spacing: single
**Formatted:** Space Before: 3 pt, After: 3 pt, Line spacing: single
**Formatted:** Space Before: 3 pt, After: 3 pt, Line spacing: single
**Formatted:** Space Before: 3 pt, After: 3 pt, Line spacing: single

C2 - Confidential Business Data

**Supplementary Table 10. Correlation between LC-MS/MS 1 hour degradation with molecular parameters**

| Property | R | P (two-tailed) | Slope | Y-intercept |
|---|---|---|---|---|
| MW | -0.097 | 0.637 | -0.189 | 318.500 |
| LD50 (μM) | 0.174 | 0.395 | 0.446 | 9.934 |
| HBA | 0.026 | 0.901 | 0.001 | 1.765 |
| HBD | -0.051 | 0.806 | -0.001 | 0.192 |
| LogP | -0.353 | 0.077 | -0.019 | 3.904 |
| *PSA | 0.380 | 0.056 | 0.189 | 46.780 |
| Rotatable bonds | -0.190 | 0.353 | -0.011 | 7.182 |
| VWR (Å) | -0.168 | 0.411 | -0.308 | 285.600 |
| BE | 0.032 | 0.878 | 0.001 | -6.325 |
| D | -0.064 | 0.757 | -0.006 | 8.880 |
| $BE_{eff}$ | -0.126 | 0.540 | -1.83E-05 | -0.021 |
| SD | -0.294 | 0.146 | -0.012 | 4.902 |
| BE(SD) | -0.301 | 0.135 | -0.011 | -3.846 |
| $BE_{eff}$(SD) | -0.334 | 0.096 | -4.45E-05 | -0.013 |

Parameters with * are significantly correlated with experimental result; Pearson Correlation

**Formatted:** Space Before: 3 pt, After: 3 pt, Line spacing: single

**Formatted:** Space Before: 3 pt, After: 3 pt, Line spacing: single

**Formatted:** Space Before: 3 pt, After: 3 pt, Line spacing: single

**Formatted:** Space Before: 3 pt, After: 3 pt, Line spacing: single

**Formatted:** Space Before: 3 pt, After: 3 pt, Line spacing: single

**Formatted:** Space Before: 3 pt, After: 3 pt, Line spacing: single

**Formatted:** Space Before: 3 pt, After: 3 pt, Line spacing: single

**Formatted:** Space Before: 3 pt, After: 3 pt, Line spacing: single

**Formatted:** Space Before: 3 pt, After: 3 pt, Line spacing: single

**Formatted:** Space Before: 3 pt, After: 3 pt, Line spacing: single

**Formatted:** Space Before: 3 pt, After: 3 pt, Line spacing: single

**Formatted:** Space Before: 3 pt, After: 3 pt, Line spacing: single

**Formatted:** Space Before: 3 pt, After: 3 pt, Line spacing: single

**Formatted:** Space Before: 3 pt, After: 3 pt, Line spacing: single

**Formatted:** Line spacing: single

C2 - Confidential Business Data

**Supplementary Table 11. Correlation between LC-MS/MS 2 hour degradation with pflour50**

| pFlour50 | R | P (two-tailed) | Slope | Y-intercept |
|---|---|---|---|---|
| *1 µM | -0.503 | 0.028 | -0.735 | 77.500 |
| *1 µM 10 min | -0.498 | 0.030 | -0.622 | 84.630 |
| 1 µM 30 min | 0.227 | 0.351 | 0.113 | 7.130 |
| *10 µM | 0.505 | 0.027 | 0.899 | -29.750 |
| *10 µM 10 min | -0.475 | 0.040 | -0.398 | 72.450 |
| *10 µM 30 min | -0.708 | 0.001 | -1.297 | 102.200 |

Parameters with * are significantly correlated with experimental result; Pearson Correlation

**Formatted:** Space Before: 3 pt, After: 3 pt, Line spacing: single

**Formatted:** Space Before: 3 pt, After: 3 pt, Line spacing: single

**Formatted:** Space Before: 3 pt, After: 3 pt, Line spacing: single

**Formatted:** Space Before: 3 pt, After: 3 pt, Line spacing: single

**Formatted:** Space Before: 3 pt, After: 3 pt, Line spacing: single

**Formatted:** Space Before: 3 pt, After: 3 pt, Line spacing: single

**Formatted:** Space Before: 3 pt, After: 3 pt, Line spacing: single

**Formatted:** Line spacing: single

C2 - Confidential Business Data

**Supplementary Table 12. Correlation between LC-MS/MS 1 hour degradation with pflour50**

| **pFlour50** | **R** | **P (two-tailed)** | **Slope** | **Y-intercept** |
|---|---|---|---|---|
| *1 µM | -0.457 | 0.028 | -0.632 | 75.790 |
| *1 µM 10 min | -0.543 | 0.007 | -0.657 | 89.940 |
| 1 µM 30 min | -0.048 | 0.827 | -0.026 | 14.160 |
| 10 µM | 0.402 | 0.057 | 0.687 | -23.290 |
| *10 µM 10 min | -0.528 | 0.010 | -0.485 | 80.460 |
| *10 µM 30 min | -0.601 | 0.002 | -1.172 | 103.800 |

Parameters with * are significantly correlated with experimental result; Pearson Correlation

**Formatted:** Line spacing: single

**Formatted:** Font: Bold

**Formatted:** Space Before: 3 pt, After: 3 pt, Line spacing: single

**Formatted:** Space Before: 3 pt, After: 3 pt, Line spacing: single

**Formatted:** Space Before: 3 pt, After: 3 pt, Line spacing: single

**Formatted:** Space Before: 3 pt, After: 3 pt, Line spacing: single

**Formatted:** Space Before: 3 pt, After: 3 pt, Line spacing: single

**Formatted:** Space Before: 3 pt, After: 3 pt, Line spacing: single

**Formatted:** Space Before: 3 pt, After: 3 pt, Line spacing: single

C2 - Confidential Business Data

**Supplementary Table 11 Correlation P-values between molecular parameters**

| | MW | LD50 | HBA | HBD | LogP | PSA | RRB | VWR | BE | D | $BE_{eff}$ | SD | BE (SD) | $BE_{eff}$ (SD) |
|---|---|---|---|---|---|---|---|---|---|---|---|---|---|---|
| MW | | NS | NS | NS | **** | NS | *** | **** | NS | NS | NS | NS | NS | *NS |
| LD50 | NS | | NS | NS | NS | NS | NS | NS | NS | NS | NS | NS | NS | NS |
| HBA | NS | NS | | NS | NS | **** | NS | NS | NS | NS | NS | NS | NS | NS |
| HBD | NS | NS | NS | | ** | NS | NS | NS | NS | NS | NS | NS | NS | NS |
| LogP | **** | NS | NS | ** | | ** | NS | **** | NS | NS | NS | NS | NS | NS |
| PSA | NS | NS | **** | NS | ** | | NS | NS | NS | NS | NS | NS | NS | NS |
| RRB | ** | NS | NS | NS | NS | NS | | **** | NS | NS | NS | NS | NS | NS |
| VWR | **** | NS | NS | NS | **** | NS | **** | | NS | NS | NS | NS | NS | NS |
| BE | NS | NS | NS | NS* | NS | NS | NS | NS | | **** | **** | **** | **** | ** |
| D | NS | NS | NS | NS | NS | NS | NS | NS | **** | | **** | **** | **** | * |
| $BE_{eff}$ | NS | NS | NS | NS | NS | NS | NS | NS | **** | **** | | **** | **** | ** |
| SD | NS | NS | NS | NS | NS | NS | NS | NS | **** | **** | **** | | **** | **** |
| BE (SD) | NS | NS | NS | NS* | NS | NS | NS | NS | **** | **** | **** | **** | | NS |
| $BE_{eff}$ (SD) | **** | NS | NS | NS | NS | NS | NS | NS | ** | * | ** | **** | NS | |

**Formatted:** Line spacing: single

**Formatted:** Space Before: 3 pt, After: 3 pt

**Formatted:** Space Before: 3 pt, After: 3 pt

**Formatted:** Space Before: 3 pt, After: 3 pt

**Formatted:** Space Before: 3 pt, After: 3 pt

**Formatted:** Space Before: 3 pt, After: 3 pt

**Formatted:** Space Before: 3 pt, After: 3 pt

**Formatted:** Space Before: 3 pt, After: 3 pt

**Formatted:** Space Before: 3 pt, After: 3 pt

**Formatted:** Space Before: 3 pt, After: 3 pt

**Formatted:** Space Before: 3 pt, After: 3 pt

**Formatted:** Space Before: 3 pt, After: 3 pt

**Formatted:** Space Before: 3 pt, After: 3 pt

**Formatted:** Space Before: 3 pt, After: 3 pt

**Formatted:** Space Before: 3 pt, After: 3 pt

**Formatted:** Line spacing: single

C2 - Confidential Business Data
